# The conserved *SEN1* DNA:RNA helicase has multiple functions during yeast meiosis

**DOI:** 10.1101/2025.04.10.648140

**Authors:** Robert Gaglione, Jonathan Caradonna, Amy J. MacQueen, Ed Luk, Nancy M. Hollingsworth

## Abstract

Diploid *Saccharomyces cerevisiae* cells undergo meiosis when they are starved of nitrogen in the presence of a non-fermentable carbon source. Nutrient starvation triggers expression of Ime1, a master regulatory protein required to activate transcription of meiotic “early genes” that mediate premeiotic S phase and prophase I processes, including recombination and chromosome synapsis. During prophase I, the highly conserved, toposomerase-like protein, Spo11, creates double strand breaks that are used to identify homologous chromosomes and generate crossovers between them. DNA:RNA hybrids are formed when an RNA molecule anneals to a complementary strand of DNA and are present at the ends of double strand breaks during prophase I of meiosis in a variety of organisms. DNA:RNA hybrids can be removed by degradation of the RNA by RNase H or by unwinding of the RNA by an essential, multi-functional DNA:RNA helicase called Sen1. Sen1 is orthologous to the mammalian Senataxin (*SETX*) helicase. Phenotypic characterization of mouse mutants lacking either Senataxin or RNase H activity exhibit male infertility and defects in double strand break repair. *SETX* is also required for meiotic sex chromosome inactivation, making it unclear whether *SETX*’s role in meiotic recombination is direct or an indirect consequence due to defects in *SETX* functions that affect transcription. Using a variety of orthogonal approaches, this work demonstrates that *SEN1* has multiple, temporally distinct roles that promote yeast meiosis. First, it enables the timely expression of *IME1*-regulated early genes. Second, it helps prevent and/or remove DNA:RNA hybrids that form during premeiotic S phase. Third, it facilitates both repair of Spo11 generated double strand breaks generated during prophase I and chromosome synapsis.

**AUTHOR SUMMARY:** DNA:RNA hybrids are unusual structures found throughout the genomes of many species, including yeast and mammals. While DNA:RNA hybrids may promote various cellular functions, persistent hybrids lead to double strand breaks, resulting in genomic instability. DNA:RNA hybrid formation and removal are therefore highly regulated by enzymes that either degrade or unwind RNA from the hybrid. Meiosis is the specialized cell division that creates haploid gametes for sexual reproduction. Previous work in yeast and mammals showed that elimination of DNA:RNA hybrids by RNase H facilitates meiotic recombination. This work demonstrates that the conserved Sen1 DNA:RNA helicase regulates the presence of DNA:RNA hybrids in three temporally distinct processes during yeast meiosis. First, *SEN1* allows for meiosis-specific genes to be expressed at the proper time to allow entry into meiosis. Second, *SEN1* prevents the accumulation of hybrids during premeiotic DNA replication. Third, *SEN1* promotes the repair of programmed meiotic double strand breaks that are necessary to form crossovers between homologous chromosomes to allow their proper segregation at the first meiotic division. Given the evolutionary conservation of Sen1 with its mammalian counterpart, Senataxin, studies of Sen1 function in yeast are likely to be informative about the regulation of DNA:RNA hybrids during humans as well.

## INTRODUCTION

DNA:RNA hybrids arise in cells when an RNA molecule base pairs with a complementary single strand of DNA. If the RNA associates with a duplex of DNA, the DNA strand of like polarity is displaced, creating a specific structure called an R-loop. It is estimated that 5-10% of the genome contains DNA:RNA hybrids [1]. DNA:RNA hybrids have beneficial effects in a variety of cellular processes, including transcriptional termination, chromatin structure, kinetochore function at metaphase I of meiosis and repression of anti-sense transcription [2–5]. However, R-loops that persist lead to the creation of double strand breaks (DSBs), resulting in genomic instability [6–10].

DNA:RNA hybrids present at the ends of DSBs may either help or hinder DSB repair by promoting resection or blocking recombinases from binding to resected single stranded (ss) ends, respectively [1, 11–17]. Furthermore, in some cases, the presence of too few or too many hybrids is deleterious, indicating that the amount of DNA:RNA hybrids must be tightly regulated [3, 17].

The importance of regulating DNA:RNA hybrids is also evident by the numerous mechanisms and proteins involved in preventing or removing them [1, 18, 19]. One way to eliminate DNA:RNA hybrids is degradation of the RNA by the conserved RNase H1 and RNase H2 enzymes [7, 20, 21]. An alternative mechanism is to unwind the RNA from the DNA using a DNA:RNA helicase such as the conserved, essential 5’-3’ yeast helicase, Sen1. Sen1 is orthologous to the mammalian helicase, Senataxin [22–25]. The Sen1 protein is multi-functional; for example, in addition to resolving R-loops, Sen1 forms a complex with Nrd1 and Nab3 to mediate transcriptional termination of unstable non-coding RNAs, as well small nucleolar RNAs. Through this activity, Sen1 influences the distribution of RNA polymerase II across the genome [22].

Given that DNA:RNA hybrids influence the repair of DSBs in vegetatively growing cells, an interesting question is whether they affect DSB repair of the programmed DSBs that initiate recombination during meiosis. Recent work looking at meiosis in mice, nematodes, and yeast defective in RNase H activity supports this idea. Mice in which RNase H1 was specifically knocked down in the germline exhibit male infertility, and spermatocytes display DSB repair defects and prophase I arrest [11]. DNA:RNA hybrids also accumulate in the nematode germline in *rnh-1.0 rnh-2* mutants [26].

Budding yeast strains containing *rnh1Δ rnh201Δ* lack RNase H activity and the resulting defects are exacerbated by deletion of *HPR1*, which disrupts the THO complex that binds to nascent messenger RNAs and assists in their maturation and nuclear export [17, 27–29]. Both *rnh1Δ rnh201Δ* and *rnh1Δ rnh201Δ hpr1Δ* diploids exhibit delayed meiotic progression and reduced spore viability [17, 30, 31].

Cytological and genomic studies in yeast using the S9.6 antibody to detect DNA:RNA hybrids in mutants lacking RNase H activity showed that hybrids form on ssDNA at the ends of DSBs generated by the meiosis-specific Spo11 protein during prophase I [17, 31, 32]. Furthermore, in both yeast and mammals lacking RNase H activity, loading of the Rad51 and Dmc1 recombinases to ssDNA ends is impaired and interhomolog recombination is reduced by the presence of DNA:RNA hybrids [11, 17]. Interestingly, experiments mapping the genomic locations of DNA:RNA hybrids in *rnh1Δ rnh201Δ* diploids showed that hybrids also accumulate during premeiotic S phase at places where there are head-on, but not co-directional, collisions between replication and transcription (called TRCs for Transcription Replication Conflicts) [31].

Given that RNase H-mediated degradation of DNA:RNA hybrids promotes meiotic DSB repair, is the same true for Senataxin/Sen1 helicase activity? *SETX*, the gene that encodes Senataxin, is required for proper mammalian meiosis, as *SETX^-/-^* mutants exhibit male infertility, as well as defects in DSB repair and crossover formation, accumulation of DNA:RNA hybrids and increased apoptosis during prophase I [33, 34]. In addition, *SETX* is required for meiotic sex chromosome inactivation (MSCI) [33, 35]. To see if *SEN1* similarly affects meiotic DSB repair in yeast, phenotypic analysis was conducted in this study using meiotic cells depleted for Sen1.

Meiosis in yeast occurs in the context of a larger process called sporulation, which results in the packaging of the four haploid genomes into gametes called spores [36]. Diploid cells are induced to sporulate when they are starved for nitrogen in the presence of a nonfermentable carbon source. These conditions result in the expression of Ime1 which, in a complex with Ume6, activates the transcription of a set of “early” meiosis-specific genes [37–41]. The early genes include those necessary for premeiotic DNA synthesis, meiotic recombination and synaptonemal complex formation.

Recombination occurs when the 5’ ends of Spo11-generated DSBs are resected, creating 3’ ssDNA tails to which the Rad51 and Dmc1 recombinases bind [42–44]. The resulting nucleoprotein filaments search the genome for homology and preferentially mediate strand invasion of homologs to form intermediates that are then processed to become either crossovers or noncrossovers [45, 46]. An intermediate step in the crossover recombination pathway triggers synapsis, the process whereby a tripartite structure called synaptonemal complex (SC) assembles between homologous chromosomes [47].Crossovers, in combination with sister chromatid cohesion, physically connect homologs, allowing them to segregate properly at Meiosis I.

The repair of DSBs is monitored by the meiotic recombination checkpoint (MRC) [48, 49]. The meiosis-specific kinase Mek1 is activated by DSBs [50, 51]. Mek1 inhibits the meiosis-specific transcription factor Ndt80 by phosphorylating its DNA binding domain [52]. As DSBs are repaired, Mek1 activity is decreased, allowing Ndt80 to bind the promoters of its target genes and activate transcription of the “middle genes” such as *CDC5*, the polo-like kinase that enables the completion of recombination and exit from prophase I, as well as genes required for Meiosis I and II and spore formation [36, 53, 54].

This work demonstrates that *SEN1* facilitates three temporally distinct processes during sporulation. First, *SEN1* promotes the timely expression of Ime1-Ume6 regulated early genes to allow entry into meiosis. Second, *SEN1* prevents the accumulation of DNA:RNA hybrids during premeiotic S phase. Third, during prophase I, *SEN*1 promotes the repair of Spo11-generated DSBs and is required for proper chromosome synapsis.

## RESULTS

### *SEN1* promotes meiotic progression, sporulation and spore viability in yeast

Because *SEN1* is essential, a meiotic depletion allele (*sen1-md*) was created by placing *SEN1* under transcriptional control of the *CLB2* promoter, which is not active during meiosis [55, 56]. Sen1 protein levels were monitored using a newly developed α-Sen1 antibody. To validate the antibody, *SEN1* and *sen1-ΔN*, (an allele that encodes an N-terminal truncation that lacks the first 1003 amino acids out of 2231) were placed under the control of the meiosis-specific *REC8* promoter and transformed into a *sen1-md* diploid [57]. Throughout the text, *sen1-md*::*PREC8-SEN1* strains are referred to simply as *PREC8-SEN1*. The diploids were then induced to undergo meiosis. Protein samples from different time points were analyzed on immunoblots probed with α-Sen1 antibodies. In the *PREC8-SEN1* diploid, a single band corresponding to Sen1’s predicted molecular weight of 253 kD was observed (S1 Fig). Furthermore, this protein was induced after four hours in Spo medium compared to vegetative cells (0 hours), consistent with *SEN1* expression being driven by the *REC8* promoter after transfer to Sporulation (Spo) medium. In the extracts from the *PREC8-sen1-ΔN* diploid, the ∼250 kD band was replaced by a less abundant protein of ∼150 kD, which is close to the expected molecular weight of the Sen1^1004-2231^ truncated fragment (135 kD), confirming the antibody’s specificity for Sen1 (S1 Fig).

Timecourses using *SEN1* and *sen1-md* diploids confirmed the depletion of the Sen1 protein during meiosis. In the wild-type (WT) diploid, Sen1 protein levels remained relatively constant throughout meiosis (Fig 1A). By contrast, the amount of Sen1 protein in vegetative cells at 0 hour in the *sen1-md* diploid was reduced relative to WT. Sen1 protein levels decreased to below the limit of detection by 3 hours in Spo medium (Fig 1A).

**Fig 1.**
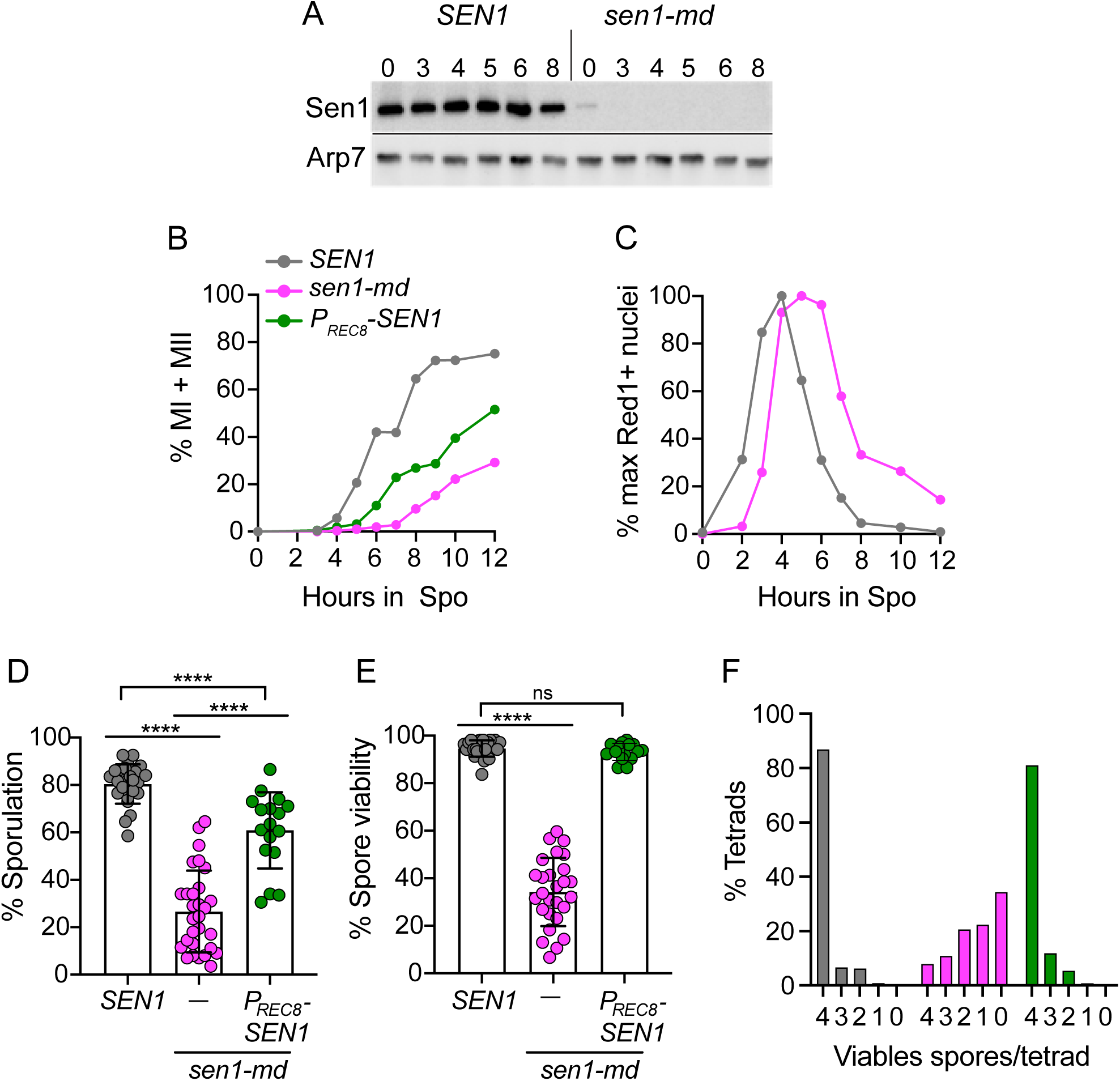
Meiotic depletion of Sen1 results in a prophase delay and reduces sporulation and spore viability. (A) Protein extracts from the indicated meiotic timepoints were generated from *SEN1* (NH716) and *sen1-md* (NH2667) diploids and probed on immunoblots with the α-Sen1 antibody. (B) Meiotic progression was followed by staining fixed cells at the indicated timepoints with DAPI and determining the frequency of MI (binucleate) and MII (tetranucleate) cells by fluorescence microscopy. 200 cells were counted at each timepoint for each replicate. *SEN1* (*n* = 7); *sen1-md* (*n* = 10); *PREC8-SEN1* (NH2667::pNH410) (*n* = 9). The averages of the biological replicates for each strain were plotted. (C) Whole cell Red1 immunofluorescence. To determine the frequency of cells in prophase I, whole cell immunofluorescence using α-Red1 antibodies was performed at the indicated timepoints in *SEN1* and *sen1-md* diploids. The % maximum Red1^+^ cells from an average of five replicates was plotted. (D) Sporulation. Cultures derived from different single colonies of the indicated strains were sporulated in liquid Spo medium and the number of asci out of 200 cells was determined by light microscopy. Each dot represents a biological replicate. *SEN1* (*n* = 25); *sen1-md* (*n* = 29); *PREC8-SEN1* (*n* = 17) (E) Spore viability. Tetrads from the cultures sporulated in Panel D were dissected to determine the frequency of viable spores. Each dot represents the spore viability of one biological replicate containing 23-28 dissected tetrads. The statistical significance of differences between the strains in Panels D and E was determined using the Mann-Whitney test (**** = *p* <0.0001). ns = not significant. (F) Distribution of viable spores in tetrads. The total number of tetrads for each strain was *SEN1* (623), *sen1-md* (728) and *PREC8-SEN1* (395).

Phenotypic characterization of the *sen1-md* diploid revealed several meiotic defects. Meiotic progression was examined by staining the DNA with a fluorescent dye and counting the number of nuclei per cell. Binucleate cells indicate completion of Meiosis I (MI), while tetra-nucleate cells indicate completion of Meiosis II (MII) (S2 Fig). On average, <30% of *sen1-md* cells progressed past MI, and those that did were delayed relative to WT (Fig 1B). In addition, both sporulation and spore viability were significantly reduced in the *sen1-md* diploid (Fig 1D, 1E).

Mutants that decrease interhomolog crossovers exhibit higher levels of inviable spores due to MI non-disjunction. As a result, the number of tetrads in which all four spores are viable goes down, with a coordinated increase in the number of tetrads with two and zero viable spores [58]. This pattern was not observed for *sen1-md* (Fig 1F).

While the number of tetrads with 4 viable spores (4:0) was greatly reduced, all other tetrad classes (3:1, 2:2, 1:3, 0:4) were increased, indicating that *sen1-md* spore lethality is unlikely due to MI chromosome mis-segregation (Fig 1F).

While the reduced amount of Sen1 protein present in vegetative *sen1-md* cells is sufficient for viability, it may not be sufficient for other *SEN1* functions (Fig 1A). This fact raises the possibility that the meiotic progression delay, reduced sporulation and decreased spore viability phenotypes result from DNA damage incurred during vegetative growth, rather than defects in meiotic processes. To test this idea, a *PREC8- SEN1* strain was examined to see if expressing *SEN1* after the initiation of meiosis meiosis is able to complement *sen1-md* meiotic phenotypes. In fact, *PREC8-SEN1* partially rescued the meiotic progression delay and sporulation defect (Fig 1B, 1D) and restored spore viability to WT levels (Fig 1E). Thus, meiosis-specific expression of *SEN1* is sufficient to reverse the meiotic defects of the *sen1-md* mutant, suggesting a requirement for Sen1 activity during normal meiosis.

A delay in meiotic progression was previously observed in the absence of RNase H activity (*rnh1Δ rnh201Δ*) where it is known that DNA:RNA hybrids form on the single-stranded tails of DSBs, thereby making them difficult to repair [17, 31]. The MRC delays exit from prophase I when a threshold number of DSBs is present. If the meiotic progression delay in *sen1-md* is also due to impaired meiotic DSB repair, then cells should take longer to exit prophase I. Prophase I cells can be distinguished using whole cell immunofluorescence with antibodies that detect Red1, a meiosis-specific axial element protein that is degraded upon exit from pachytene [54, 59, 60]. This analysis found that both entry into and exit from prophase I were affected in the *sen1-md* diploid, indicating there are two separate components to the meiotic progression delay. First, while the upward slopes representing progression to a mid-prophase I stage (where Red1 is maximally abundant in pachytene) were similar between *SEN1* and *sen1-md*, the *sen1-md* cells reached a peak level of maximal mid-prophase I nuclei one hour later than WT (Fig 1C). Second, the *sen1-md* downward slope due to exit from prophase I was shallower than WT and almost 15% of the cells remained in prophase I at 12 hours (Fig 1C). This result indicates that *SEN1* is required for timely progression into and through prophase I.

### *SEN1* promotes expression of *IME1/UME6* regulated genes

*REC8* is an early meiotic gene whose transcription is activated by the Ime1- Ume6 complex [37, 38, 41, 61]. One explanation for why *PREC8-SEN1* only partially rescues the *sen1-md* meiotic progression delay is that *SEN1* promotes the timely transcription of genes regulated by Ime1-Ume6. As a first test of this hypothesis, the timing of expression for several early meiosis-specific proteins encoded was examined.

*HOP1*, *RED1* and *REC8* encode meiosis-specific components of the axial elements formed along sister chromatids [59, 61–63]. *MEK1/MRE4* encodes a meiosis-specific kinase that regulates many different steps of meiotic recombination, as well as the MRC [52, 64–70]. Hed1 is a meiosis-specific protein that binds to Rad51 to inhibit its strand exchange activity during prophase I [71, 72]. Mek1 phosphorylation of Hed1 threonine 40 stabilizes the protein and can be used as a marker for Mek1 kinase activity [53, 73].

In the WT diploid, Rec8 protein was first observed after 2 hours in Spo medium, its level peaked at 4 hours and was nearly undetectable by 8 hours (S1B Fig). By 4 hours, most of the cells were in prophase I, as indicated by peak levels of Red1 and meiotic DSBs [indirectly indicated by Hop1 phosphorylation (pHop1)] (Fig 1C, S1B Fig) [74]. Mek1 activity, indicated by phosphorylated Hed1(pHed1), also peaked at four hours, in keeping with the larger amount of phosphorylated Hop1 (S1B Fig)[73]. Red1 was nearly gone by 6 hours consistent with cells progressing into MI (Fig 1B, 1C and S1B Fig). In contrast, all 5 proteins in the *sen1-md* diploid peaked later than 4 hours and were still present at 10 hours, consistent with a portion of the cells being delayed in prophase I. Notably, the Rec8 protein from the *PREC8-SEN1* diploid exhibited similar expression kinetics as in *sen1-md* (S1B Fig).

To more directly test the hypothesis that *SEN1* promotes early gene transcription, RNA sequencing analysis was performed using cells taken at different times after transfer to Spo medium. As was previously observed, meiotic progression in the *sen1- md* diploid was delayed in this time course relative to WT (Fig 2A). Genes were grouped by k-means based on RNA expression patterns in the *SEN1* diploid. The heat map for the WT strain exhibited gene clusters that agreed well with the literature (Fig 2B) [37, 41]. When a subset of early genes from the WT diploid known to function in prophase I was examined, their expression in the WT diploid nearly uniformly peaked at 3 to 4 hours after transfer to Spo medium (Fig 2C). In contrast, expression of the same set of genes in the *sen1-md* mutant was delayed, with broader peaks occurring between 4 and 6 hours. The delay in early gene transcription correlated with the later appearance of *IME1* transcripts, the master regulator for early gene transcription, in the *sen1-md* strain compared to WT (Fig 2C). As expected, the delay in early gene transcription coordinately delayed the expression of *NDT80*-regulated middle genes (Fig 2B, 2C). We propose that meiotic progression proceeds more slowly in *sen1-md* for two reasons: (1) a delay in the onset of early gene transcription, and (2) activation of the MRC due to the presence of unrepaired DSBs. This hypothesis would explain the partial rescue of the meiotic progression defect in *PREC8-SEN1*. Ectopic expression of *SEN1* during meiosis using *PREC8-SEN1* allows DSB repair, thereby deactivating the MRC and allowing cells to progress faster than *sen1-md*. However, because *SEN1* transcripts produced using the *REC8* promoter appeared too late to fix the early gene transcription defect, the cells still exhibited a delay in progression through the early stages of meiosis.

**Fig 2.**
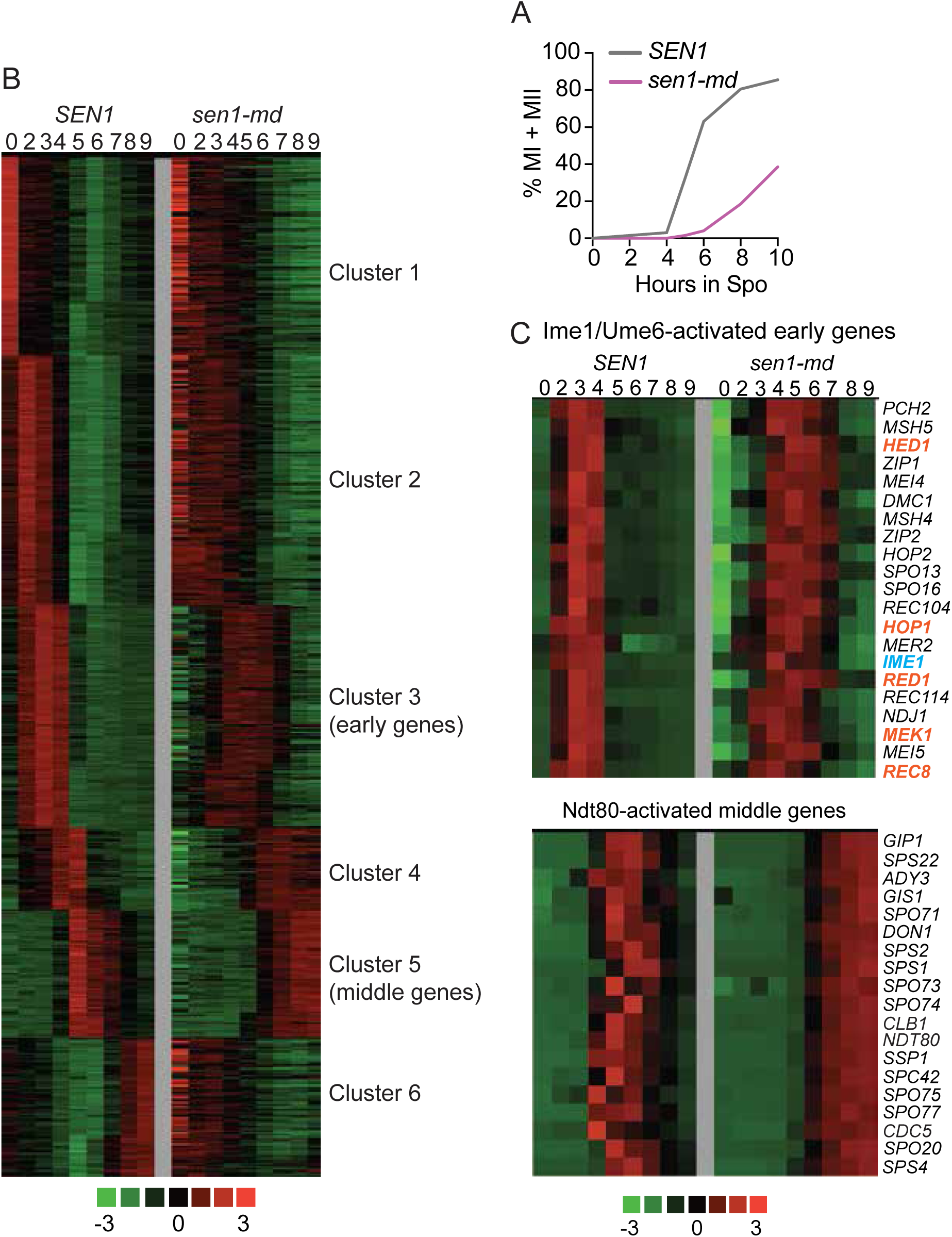
RNA sequencing analysis of *SEN1* and *sen1-md* diploids during meiosis. *SEN1* (NH2473) and *sen1-md* (NH2520) were transferred to Spo medium and cells were taken at various timepoints. (A) Meiotic progression. Fixed cells were stained with DAPI and examined by fluorescence microscopy for the presence of MI and MII cells. (B) RNA-seq analysis. mRNA was isolated at the indicated timepoints from the timecourse shown in Panel A. Numbers above each column indicate hours after transfer to Spo medium. The *z* value for each timepoint (n) was calculated as [(FPKMn- FPKMave)/SD], where FPKM = fragments per kilobase per million, SD = standard deviation and ave = average. Genes were sorted based on the RNA expression patterns in the *SEN1* diploid by clustering analysis (k-means) using Cluster 3.0. The heat map was generated using Java TreeView. Green indicates *z* values < 0, while red indicates *z* values > 0. (C) A subset of previously identified *IME1/UME6*-dependent early genes and *NDT80*-dependent middle genes were selected and clustered as in Panel B. Gene names in orange encode proteins that were examined in S1b Fig, while *IME1* is written in blue.

### The *sen1-md* delay in exiting prophase I is due to the meiotic recombination checkpoint

The MRC is controlled by the meiosis-specific Mek1 kinase [49, 52, 75]. Mek1 is activated by the presence of DSBs [50, 51, 76]. When DSB levels are high, Mek1 phosphorylates and inhibits the meiosis-specific transcription factor, Ndt80, that is required for the expression of >300 genes, including *CDC5* and *CLB1*, which are involved in completing recombination and prophase I exit [52–54, 77]. Mek1 interacts with Ndt80 through a five-amino acid sequence in the transcription factor and deletion of that sequence abrogates the MRC [52]. The *NDT80* allele containing this five-codon deletion is referred to as *NDT80-mid* (Mek1 Interaction Defective). *NDT80-mid* specifically disrupts the MRC without affecting Mek1 kinase activity and cells progress through meiosis with the same kinetics as WT [60]. Therefore *NDT80-mid* is a useful tool for determining whether the MRC is triggered by a particular mutant.

The MRC prevents or delays cells from exiting prophase I [49, 60, 78]. This phenotype can be indirectly examined by looking at the frequency of cells that proceed through MI to become binucleate. WT MI cells peaked at ∼13% after only 5 hours in Spo medium (Fig 3A). In contrast only ∼7% of the *sen1-md* cells had completed MI after 12 hours (Fig 3A). The onset of MI occurred later than WT in both *NDT80-mid sen1-md* and *sen1-md* due to the delay in early gene transcription. However, *NDT80-mid sen1- md* cells completed MI faster than *sen1-md*, peaking with ∼18% cells at 8 hours (Fig 3A). The fact that the MRC is responsible for the prophase I arrest/delay observed when Sen1 is meiotically depleted indicates that *SEN1* is required for timely DSB repair.

**Fig 3.**
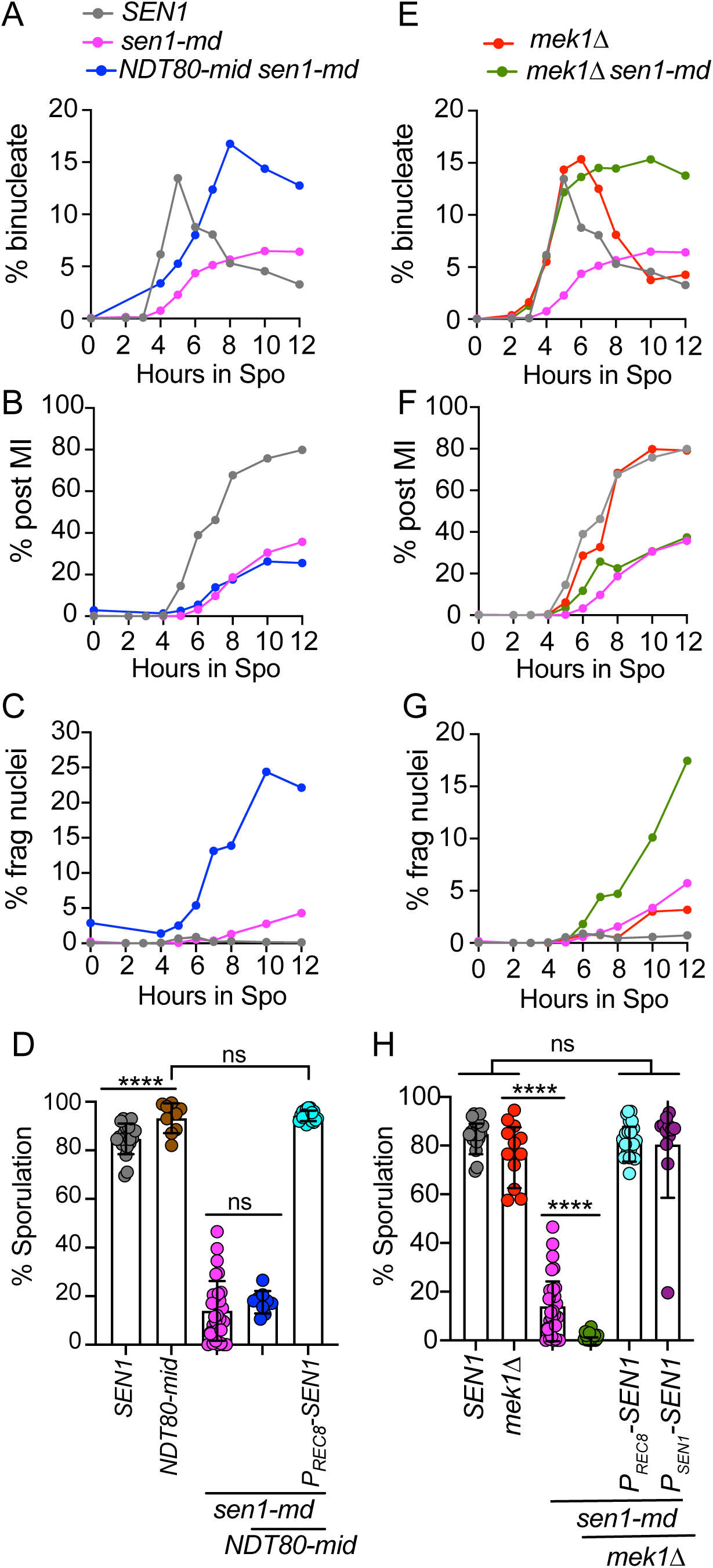
Meiotic progression and sporulation of *sen1-md* diploids containing *NDT80-mid* or *mek1Δ*. *SEN1* (*n* = 13), *sen1-md* (*n* =20), *NDT80-mid sen1-md* (NH2667::pNH317^2^) (*n* = 4), *mek1Δ* (NH729) (*n* = 6), and *mek1Δ sen1-md* (NH2669) (*n* = 10) diploids were induced to undergo meiosis and the average frequency of the indicated cell types was determined at various time points as described in Fig 1B. (A, E) binucleate cells that have completed MI; (B, F) Post-MI contains both MII cells (tetranucleate) and cells with fragmented nuclei (>4 DAPI foci). (C, G) Cells with fragmented nuclei. (D,H) Different colonies from the diploids used in strains used in Panels A and E, as well as *NDT80-mid PREC8-SEN1* (NH2667::pNH410::pNH317^2^), *mek1Δ PREC8-SEN1* (NH2669::pNH410), and *mek1Δ PSEN1-SEN1* (NH2669::pBG28), were sporulated at 30°C on solid medium. Each dot represents a biological replicate for which the percentage of sporulated cells was determined for 200 cells. The *SEN1* and *sen1-md* data are repeated in all of the graphs. Error bars indicate the means and standard deviations. Statistical significance of differences between strains was determined using the Mann-Whitney test (* = *p* <0.02; ** = *p* <0.002; *** = *p* <0.001; **** = *p* <0.0001).

Although *NDT80-mid* allows *sen1-md* cells to exit prophase I, it does not rescue the sporulation defect (Fig 3D). This may be because allowing cells to proceed through the meiotic divisions with unrepaired DSBs resulted in fragmented nuclei, as indicated by the presence of more than 4 DAPI staining foci, similar to what is observed when the MRC is inactivated in *dmc1Δ* or *sae2Δ* diploids (S2 Fig)[78, 79]. Cells containing either four nuclei (i.e complete MII) or fragmented nuclei were classified as “post-MI”. Greater than 80% of WT cells completed MII by 12 hours with <5% showing fragmented nuclei (Fig 3B, 3C). In contrast, only ∼25% of *sen1-md* and *NDT80-mid sen1-md* cells proceeded past MI (Fig 3B). A major difference between these two strains, however, was that almost all the *NDT80-mid sen1-md* post-MI cells contained fragmented nuclei, compared to <5% of *sen1-md* cells (Fig 3C). Therefore, the prophase I delay created by the MRC in the *sen1-md* diploid provided time for most DSBs to be repaired.

*MEK1* controls not only the MRC, but also interhomolog bias and inhibition of Rad51 strand exchange activity [67, 69, 73]. For example, when *MEK1* is inactive in a *dmc1Δ* mutant, cells progress through meiosis with intact chromosomes because Rad51 repairs DSBs using sister chromatids, thereby removing the signal that activates the MRC [49, 74]. In contrast, if DSBs are unrepairable using either homologs or sister chromatids as templates (eg. *sae2Δ* or *rad50S*), cells progress through meiosis with fragmented nuclei in the absence of the MRC [49, 79]. Similar to *NDT80-mid*, combining *mek1Δ* with *sen1-md* resulted in faster entry into MI compared to *sen1-md* with a reduced number of post-MI cells (Fig 3E, 3F). The increased frequency of fragmented nuclei in the *mek1Δ sen1-md* diploid suggests that DSBs persisted because they were unrepairable (Fig 3G). The sporulation defects of *NDT80-mid sen1-md* and *mek1Δ sen1-md* were rescued by *PREC8-SEN1* to the same extent as the constitutively expressed *PSEN1-SEN1*, consistent with a need for *SEN1* during prophase I to allow DSB repair (Fig 3D, 3H).

### *SEN1* facilitates meiotic recombination at the *HIS4LEU2* DSB hotspot

To determine directly whether *SEN1* promotes meiotic recombination, physical analyses using the *HIS4LEU2* hotspot were performed. This well characterized hotspot has asymmetric XhoI sites that give rise to distinct bands corresponding to DSBs and crossovers (COs) that can be detected using Southern blots (Fig 4A) [80]. In addition, there is an NgoMIV site on one homolog adjacent to the region where Spo11 makes DSBs. Digestion with both XhoI and NgoMIV therefore also allows detection of noncrossovers (NCOs) resulting from gene conversion events involving the NgoMIV site [75].

**Fig 4.**
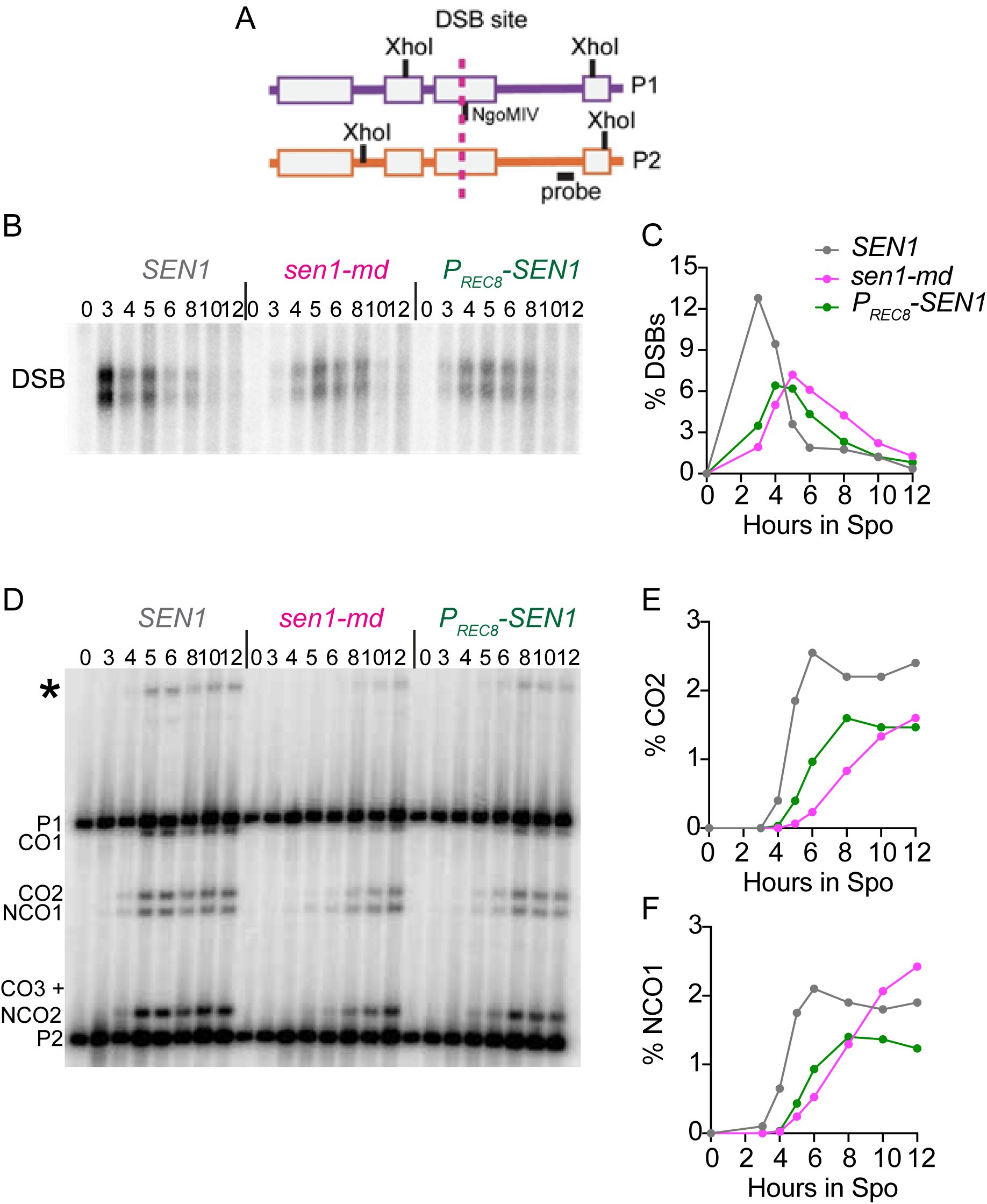
Physical analysis of DSBs, crossover and noncrossover formation during meiosis in *sen1-md* and *PREC8-SEN1* diploids. Time courses of *SEN1* (*n* = 2), *sen1- md* (*n* = 3) and *PREC8-SEN1* (*n* = 3) diploids were performed and genomic DNA isolated from cells at the indicated timepoints. (A) Schematic of the *HIS4LEU2* hotspot. The homologs, P1 and P2, have XhoI sites located at asymmetric positions. The dotted vertical line indicates where Spo11 makes DSBs. The NgoMIV site is adjacent to the DSB site on P1. The black box indicates the sequence used as a probe in Southern blot analysis. (B) DSBs: Genomic DNA was digested with XhoI to detect Spo11-generated breaks. The XhoI sites on the two homologs are slightly offset, resulting in two DSB fragments that undergo variable amounts of resection. (C) Average DSB values for the different replicates normalized to the total amount of DNA. (D) CO and NCO: Genomic DNA was digested with XhoI and NgoMIV. CO1 and CO2 indicate crossover bands, while NCO1 and NCO2 represent noncrossover bands. (E) % average CO2 values. (F) % average NCO1 values. The asterisk indicates a band containing a flanking XhoI site gene conversion.

In the WT diploid, DSBs peaked at 3 hours after transfer to Spo medium and then rapidly disappeared (Fig 4B, 4C). In contrast, DSB formation was delayed in both *sen1-md* and *PREC8-SEN1*. This was expected as transcription of most of the genes required for making DSBs (eg. *SPO11*, *REC114*, *REC102* and *REC104*) are dependent on Ime1/Ume6. DSBs also persisted longer in *sen1-md* than in either *SEN1* or *PREC8- SEN1* and were still detectable after 12 hours (Fig 4C). CO2 and NCO1 fragments that can be unambiguously identified on the XhoI/NgoMIV gel were quantified (Fig 4D). In the WT strain, CO formation plateaued by 6 hours. In contrast, CO formation in *sen1-md* was delayed and reduced by 25% at 12 hours, but still had not reached a plateau (Fig 4D, 4E). The kinetics of CO formation in the *PREC8-SEN1* strain was intermediate between WT and *sen1-md*, as expected if the *sen1-md* early gene transcription delay was coupled to more efficient DSB repair by *SEN1*. However, for reasons that are unclear, the level of COs in the *PREC8-SEN1* strain resembled that of *sen1-md* and did not reach WT levels.

A different phenotype was observed for NCOs in the *sen1-md* diploid. Instead of being decreased, there were more NCOs at 12 hours compared to WT (Fig 4D, 4F).

This result could be because prophase I length is longer in many of the *sen1-md* cells, which delays the expression of the polo-like kinase, *CDC5*, required for resolution of the double Holliday junctions into COs [54]. Furthermore, cells arrested in pachytene continue to make DSBs and produce NCOs by synthesis-dependent strand annealing [81–83]. The idea that the increase in NCO formation in *sen1-md* is due to a delay in prophase I exit is supported by the observation that the *PREC8-SEN1* diploid, in which cells were able to progress, NCOs plateaued by 8 hours (Fig 4D, 4F).

### *SEN1* promotes chromosome synapsis during meiosis

In budding yeast, meiotic DSB repair by a crossover specific pathway containing a functionally diverse set of “ZMM” proteins is required for SC formation [84–89]. The SC is a tripartite supramolecular chromosomal complex that initiates assembly from discrete chromosome sites associated with recombination (as well as from centromeres) and extends along the full lengths of aligned homologous chromosome axes [90, 91]. SC formation in yeast is completely dependent on DSB formation and early processing. Diploids containing *spo11Δ*, *sae2Δ* or *rad50S* mutants exhibit only aggregates of SC structural proteins (polycomplexes) instead of linear SC structures, while the DSB repair defects of *dmc1Δ* or *msh4Δ* are associated with an overall dcrease and delay in SC formation, rather than a complete abrogation of SC [92–98].

Diploids homozygous for *ndt80Δ* arrest in pachynema, the stage of prophase I when chromosomes are fully synapsed [99]. The ability of *ndt80Δ sen1-md* diploids to assemble SCs was assessed by examining the distribution of SC structural components Zip1 (an SC transverse filament protein) and Gmc2 (part of the SC central element) on surface-spread meiotic chromosomes that displayed maximal Red1 accumulation after 5 hours in Spo medium [100, 101]. At this timepoint, 83% (n=103) of the *ndt80Δ* control nuclei exhibited full-length or nearly full-length linear structures of Gmc2 throughout the entirety of the mid-prophase nucleus, while the remaining spread chromosomes showed shorter linear assemblies of Gmc2 (Fig. 5A, 5B). By contrast, only 4% (n=103) of the surface-spread nuclei from the *ndt80Δ sen1-md* strain exhibited full-length linear Gmc2 structures, even though most spreads displayed Red1 accumulation on chromosome axes to the same extent as the *ndt80Δ* control. The remaining nuclei were nearly evenly divided between those that had only Gmc2 foci (50%) and those with shorter linear Gmc2 assemblies (46%). In some *ndt80Δ sen1-md* cells, the Gmc2 linear assemblies detected on chromatin co-localized with Zip1, indicating that these structures correspond to mature (albeit partial) SCs (Figure 5A). Polycomplexes containing Gmc2 were observed in 26% of the *ndt80Δ* spreads; this number was increased to 49% in the *ndt80Δ sen1-md* diploid (S1 Data). *SEN1* therefore plays an important role in promoting chromosome synapsis during yeast meiosis.

**Fig 5.**
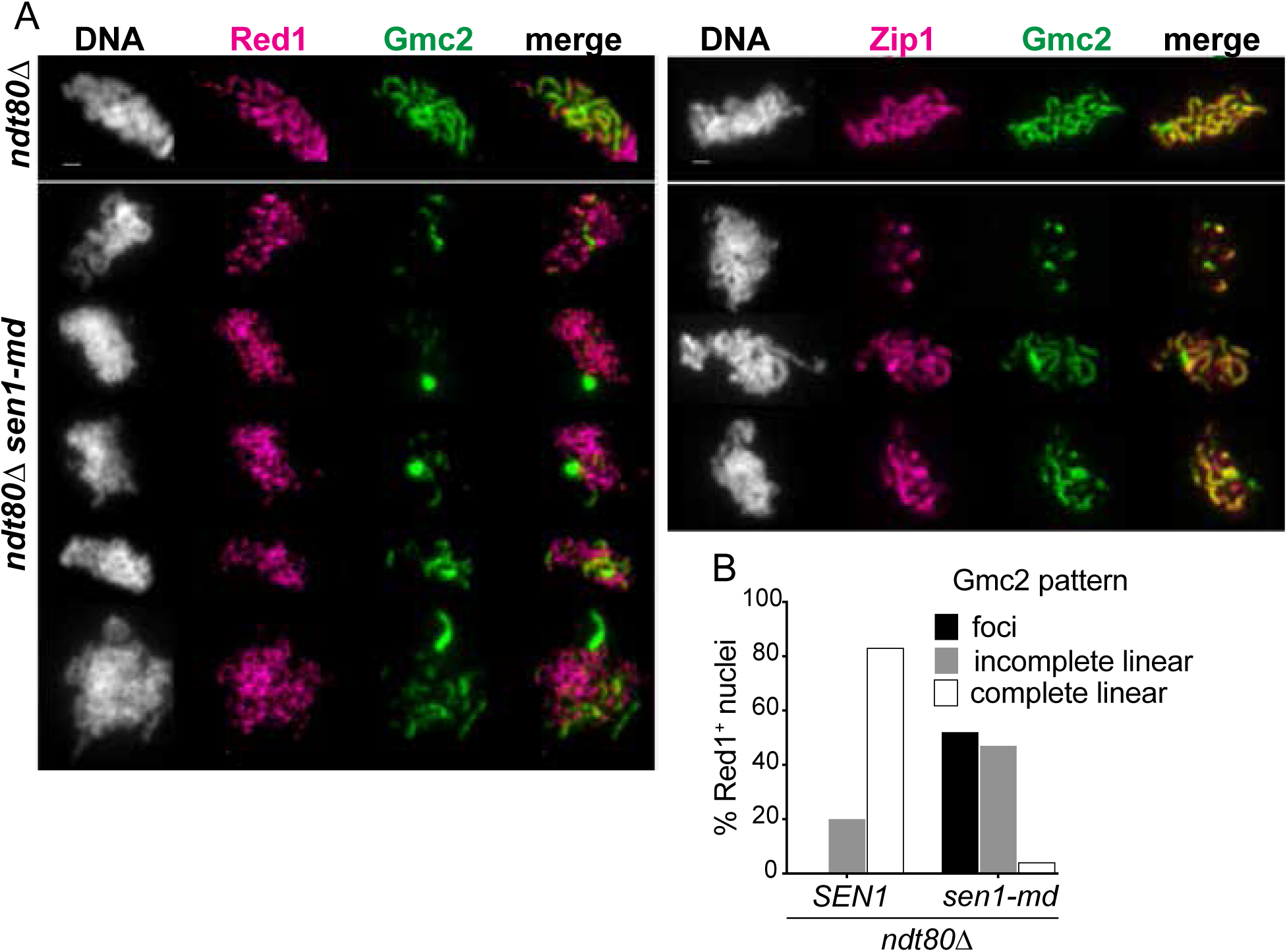
SC assembly in *ndt80Δ* and *ndt80Δ sen1-md* diploids The *ndt80Δ* (NH2188) and *ndt80Δ sen1-md* (AM6697) diploids were transferred to Spo medium for 5 hours at which time cells were processed to make chromosome spreads. (A) Immunofluorescence on surface-spread meiotic chromosomes to detect chromosome axes (Red) and the central element (Gmc2). Each row corresponds to a different surface-spread nucleus, with the genotypes indicated at left. DNA is labeled with DAPI (white, first column), anti-Red1 (magenta, second column), anti-Gmc2 (green) or anti-Zip1 (magenta, second panel). The last column shows the merged image. Bar, 1 μm. (B) Histogram of the fraction of total nuclei examined (*n*=103 for each genotype) with complete SC (full linear structures of Gmc2 that extend throughout the full spread nucleus), incomplete SC (one or more shorter linear structures of Gmc2 that only partially extend throughout the surface-spread nucleus), or only foci of Gmc2. Note that only those surface-spread nuclei with maximal Red1 accumulation on chromosome axes were included in this analysis.

### *SEN1* prevents formation of *SPO11*-independent DSBs during meiosis

If Spo11 generated meiotic DSBs are solely responsible for the MRC-mediated prophase I delay in the *sen1-md* diploid, then deleting *SPO11* should rescue this delay. In fact, the *sen1-md spo11Δ* diploid progressed faster through MI than *sen1-md*, indicating that the MRC was no longer being triggered (Fig 6A). However, these chromosomes were predicted to be intact and therefore enter MII efficiently as is true for *spo11Δ*, but this was not the case (Fig 6B). Instead, *sen1-md spo11Δ* exhibited a reduced number of post-MII cells, an increase in cells with fragmented nuclei, and decreased sporulation (Fig 6B, 6C, 6D). The presence of fragmented nuclei shows that DSBs were present in the *sen1-md spo11Δ* cells that were not directly created by Spo11.

**Fig 6.**
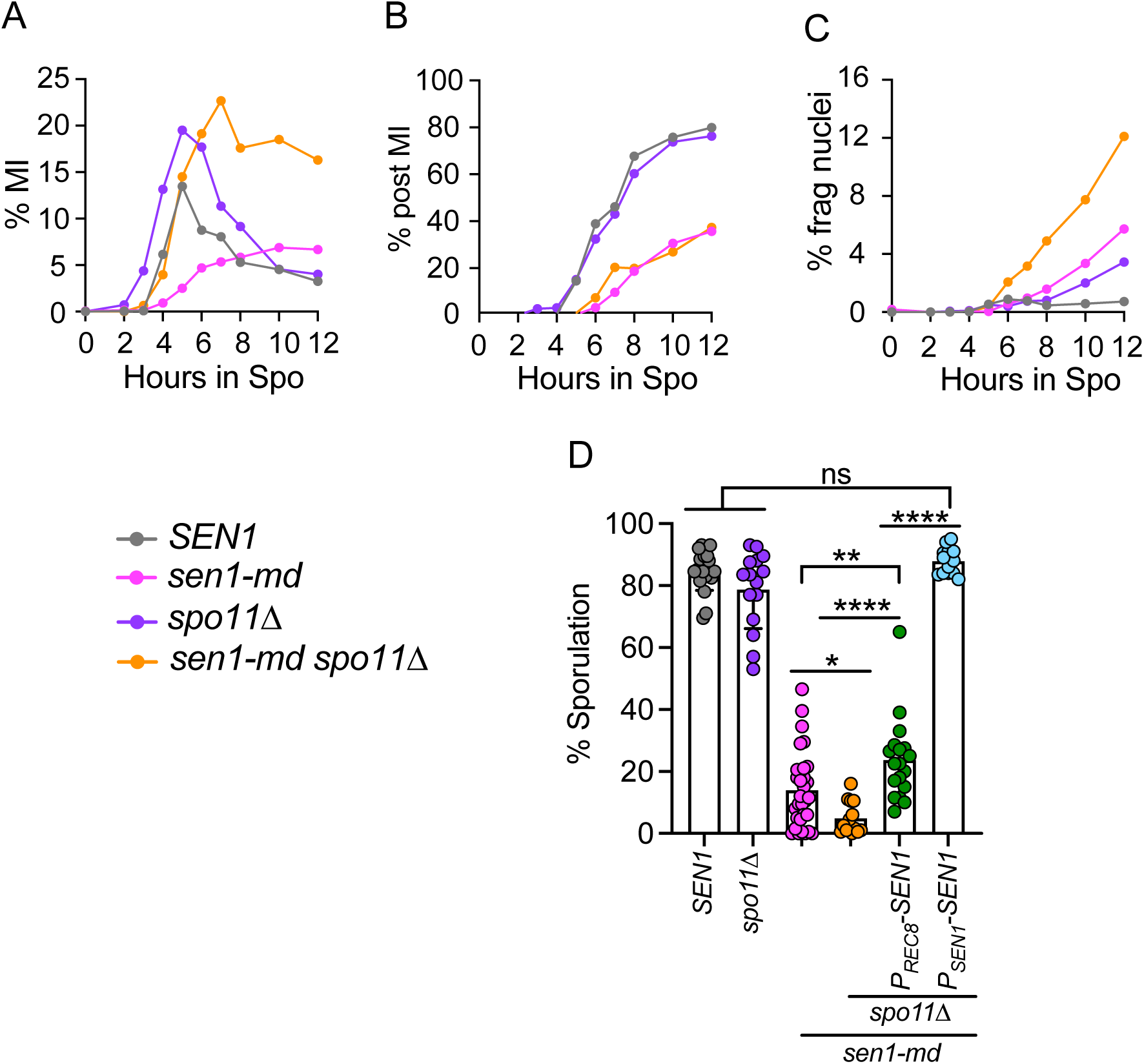
Meiotic progression and sporulation of *sen1-md* diploids containing *spo11Δ. SEN1*, *sen1-md*, *spo11Δ* (NH1055, *n* = 10) and *spo11Δ sen1-md* (NH2689, *n* = 15) diploids were induced to undergo meiosis and the average frequency of the indicated cell types was determined at various time points as described in Fig 3. The *SEN1* and *sen1-md* data for panels A-C are the same at Fig 3. (A) Cells that have completed MI; (B) Post-MI contains both MII cells and cells with fragmented nuclei (>4 DAPI foci). (C) Cells with fragmented nuclei. (D) Different colonies from the diploids used in Panels A-C, *spo11Δ PREC8-SEN1* (NH2689::pNH410) and *spo11Δ PSEN1-SEN1* (NH2689::pBG28) were sporulated at 30°C on solid medium. Each dot represents a biological replicate for which the frequency of sporulated cells was determined for 200 cells. The data for WT and *sen1-md* from Fig 3 are repeated in all of the graphs. Error bars indicate the means and standard deviations. Statistical significance of differences between strains was determined using the Mann-Whitney test (* = *p* <0.02; ** = *p* <0.002; *** = *p* <0.001; **** = *p* <0.0001).

In contrast to *NDT80-mid sen1-md* and *mek1Δ sen1-md*, *PREC8-SEN1* only weakly complemented the sporulation defect of *sen1-md spo11Δ* (Fig 6D). However, the *sen1-md spo11Δ* sporulation defect was completely rescued when *SEN1* was constitutively expressed using *PSEN1-SEN1*. This difference indicates that the timing of *SEN1* transcription matters in the absence of *SPO11* and argues that DSBs present in *sen1-md spo11Δ* are temporally separable and occur earlier than the Spo11 generated DSBs in *sen1-md*.

### *SEN1* prevents accumulation of DNA:RNA hybrids during premeiotic S phase and prophase I

The *sen1-md* meiotic phenotypes mirror those of *rnh1Δ rnh201Δ* and *rnh1Δ rnh201Δ hpr1Δ* diploids, suggesting they similarly arise from the failure to prevent DNA:RNA hybrids from accumulating during meiosis (Fig 1)[17, 31]. To test this idea, the frequency of DNA:RNA hybrids (some of which may be R-loops) was compared between *SEN1* and *sen1-md* at different times after transfer to Spo medium. DNA:RNA hybrids were detected as foci on chromosome spreads using the S9.6 antibody (Fig 7A)[32]. The *rnh1Δ rnh201Δ hpr1Δ* diploid was used as a positive control as it has previously been shown that vegetative cells with this genotype exhibit numerous DNA:RNA hybrid foci using this antibody (Fig 7A)[17]. To further validate the assay, chromosome spreads from *rnh1Δ rnh201Δ hpr1Δ* cells collected 9 hours after transfer to Spo medium were treated with RNase H prior to antibody staining. This treatment led to a significant reduction of S9.6 foci, confirming that they are DNA:RNA hybrids (Fig 7B).

**Fig 7.**
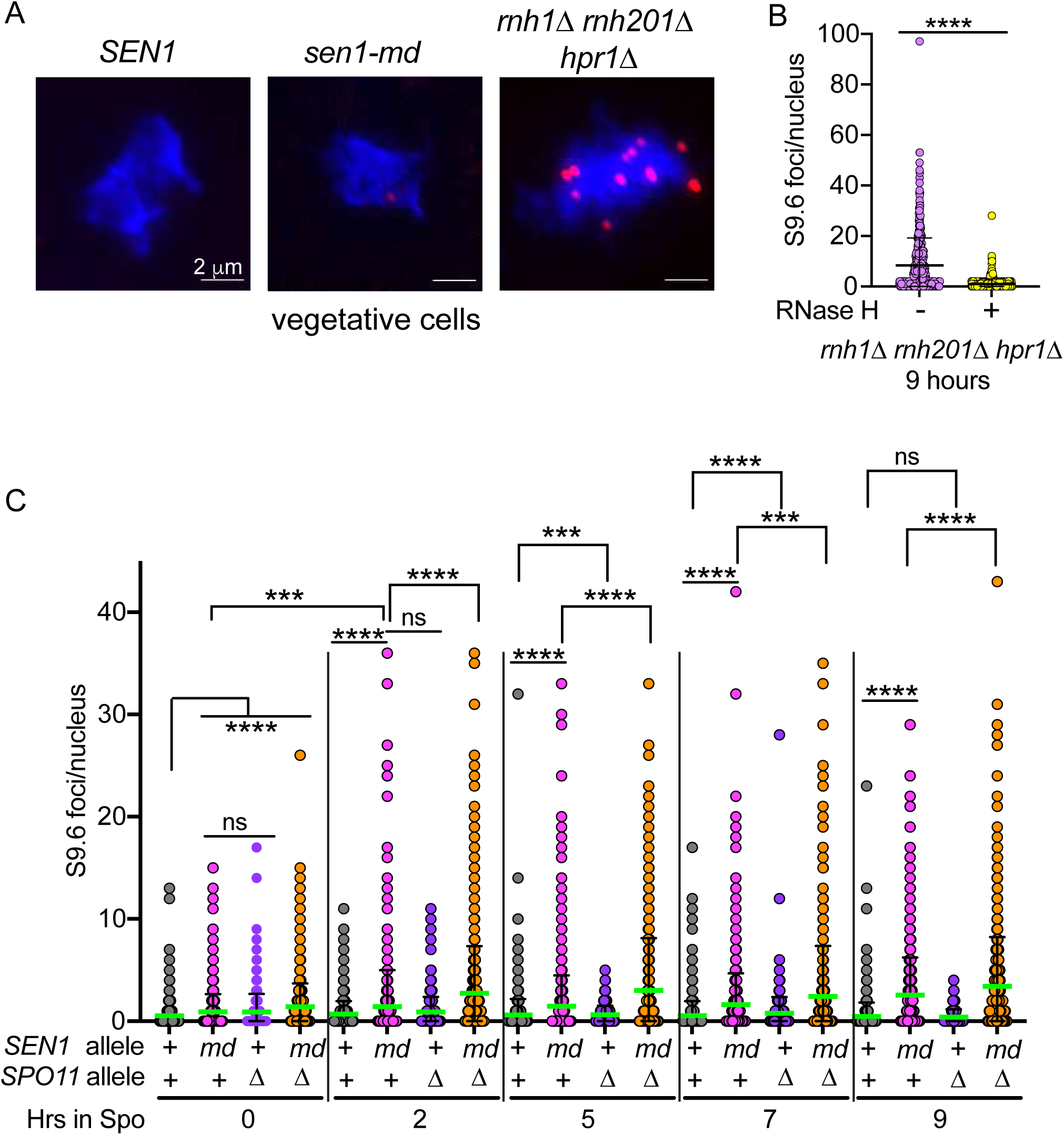
*SEN1* decreases DNA:RNA hybrids during meiosis. (A) Chromosome spreads were prepared from vegetatively growing *SEN1*, *sen1-md* and *rnh1Δ rnh201Δ hpr1Δ* (LZY2919) diploids. The DNA was stained with DAPI (blue) and DNA:RNA hybrids were detected by immunofluorescence (red foci) using the S9.6 antibody. (B) Chromosome spreads from *rnh1Δ rnh201Δ hpr1Δ* cells at the 9 hr meiotic timepoint were treated with either bovine serum albumin (-RNase H) or RNase H (+RNase H) and then stained with the S9.6 antibody (*n* = 2). (C) DNA:RNA hybrid foci were detected on chromosome spreads from *SEN1* (*n* = 4 or 5 depending on the timepoint), *sen1-md* (*n* = 4, 5 or 6), *spo11Δ* (*n* = 3) and *sen1-md spo11Δ* (*n* = 3) at the indicated timepoints using the S9.6 antibody. *n* varies for some strains depending upon the timepoint. Two hundred spreads were examined for each biological replicate. Green bars indicate the mean values. Statistical significance was determined using the Mann-Whitney test (* = *p* <0.02; ** = *p* <0.002; *** = *p* <0.001; **** = *p* <0.0001).

The *sen1-md*, *spo11Δ* and *sen1-md spo11Δ* diploids all exhibited a statistically significant increase in S9.6 foci at the 0 hour timepoint compared to WT (Fig 7C). This suggests that *sen1-md* allele is indeed hypomorphic in vegetative cells, that is, while viable, the reduced amount of Sen1 protein does have an effect. Given that *SPO11* is a meiosis-specific gene that is not expressed in vegetative cells, it is unclear why the DNA:RNA hybrid signal was elevated in the *spo11Δ* diploid at 0 hours [102]. However, while the number of S9.6 foci in *spo11Δ* remained constant until decreasing at 9 hours, the frequency of DNA:RNA hybrids increased for both *sen1-md* and *sen1-md spo11Δ* diploids starting at 2 hours and remained high up to the 9 hour timepoint (Fig 7C).

Furthermore, *spo11Δ* exacerbated the *sen1-md* phenotype, as more DNA:RNA hybrid foci were observed in the double mutant compared to *sen1-md* alone starting with the 2 hour timepoint. We conclude that during yeast meiosis, the amount of DNA:RNA hybrids is regulated by Sen1 as well as by RNase H activity.

## DISCUSSION

### *SEN1* is a regulator of meiotic early gene expression

In the wild, budding yeast cells are induced to undergo sporulation when they are starved for nutrients such as nitrogen and glucose. As a result, the haploid genomes produced by meiosis are packaged into spores that remain quiescent until nutrients return. Spores are resistant to a variety of stresses, including enzymes present in the fruit fly digestive tract, allowing yeast spores to be dispersed to new locations after being eaten [103]. Entry into meiosis is controlled by Ime1, a master transcriptional regulator that activates transcription of the “early genes” required for premeiotic S phase and prophase I [39, 104]. Because the *sen1-md* mutant delayed the expression of all the early genes as well as *IME1*, we propose that the reduced amount of Sen1 protein generated by the *CLB2* promoter delays entry into meiosis by affecting transcription of *IME1* (although alternative possibilities cannot be ruled out).

The regulation of *IME1* expression is complex, as it requires integration of signals relating to both mating type and nutrients (reviewed in [104, 105]. The promoter of *IME1* is approximately 2 kb in length [106, 107]. Within the first 1.1 kb are multiple regulatory sites including ones that repress *IME1* transcription when glucose is present or activate transcription when cells are exposed to acetate [107]. In haploid cells, this region of the *IME1* promoter exists in a repressive chromatin state due to transcription of a long noncoding RNA called *IRT1* that begins 1.4 kb upstream of the *IME1* gene [108] (Fig 8). This repressive chromatin prevents transcriptional activators such as Pog1 from binding to the *IME1* promoter [108]. *RME1* was originally identified as a repressor of meiosis [109]. Rme1 binds to two sites in the *IME1* promoter upstream of *IRT1* and its mechanism of repression is to activate transcription of *IRT1*, thereby creating the repressive chromatin upstream of *IME1* (Fig 8)[108]. Transcription of *RME1* itself is repressed in diploids by the a1/α2 repressor encoded by *MAT***a** and *MAT*α, thereby preventing transcription of *IRT1* and allowing expression of *IME1* when diploid cells are in Spo medium (Fig 8). In some strain backgrounds (but not SK1), the repression of *RME1* is leaky, thereby allowing a low level of *IRT1* transcription that reduces the ability of these strains to sporulate [108]. Transcription of a second long non-coding RNA called *IRT2* located immediately upstream of *IRT1* prevents Rme1 from binding to its sites in the *IME1* promoter, thereby, ensuring that no *IRT1* transcripts are made (Fig 8)([110]. *IRT2* transcription is activated by a complex formed between Ume6 and Ime1 [110].

**Fig 8.**
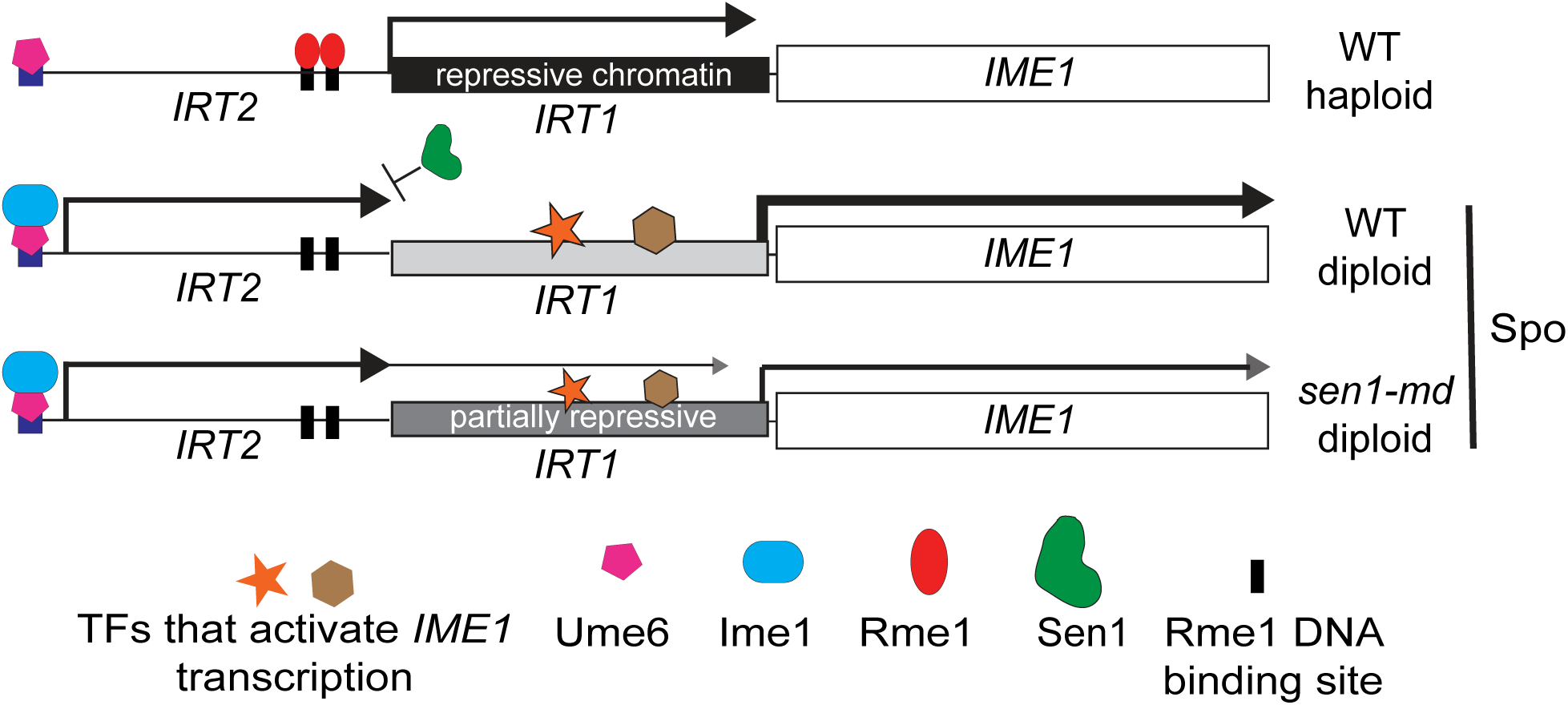
A model for *SEN1*’s role in regulating *IME1* transcription In haploid cells, *RME1*-activated transcription of the long non-coding *IRT1* RNA establishes a “repressive chromatin state” in the *IME1* promoter that prevents transcription factors from binding, resulting in a low level of *IME1* transcription. In diploid cells, the transcription of the *RME1* gene is prevented by the a1/α2 repressor, thereby preventing *IRT1* transcription [108]. Under sporulation conditions (Spo), transcription factors able to bind to the *IME1* promoter and the gene is transcribed. Ime1 then binds to Ume6 upstream of *IRT1* to activate transcription of a second long non-coding RNA called *IRT2* that further prevents Rme1 from activating *IRT1* expression [110]. We propose that *SEN1* functions in transcriptional termination of *IRT2* RNAs to prevent transcription of the *IRT1* region of the *IME1* promoter and that transcriptional termination is less efficient in the *sen1-md* diploid due to the reduced amount of Sen1 protein. As result, some *IRT2* trans cripts extend through the *IRT1* region, decreasing the amount of *IME1* transcription.

How might *SEN1* affect *IME1* regulation? The essential function of *SEN1* is the termination of RNA polymerase II non-coding RNAs as part of a complex with Nrd1 and Nab3 (the NNS complex) [111–114]. When NNS-mediated transcription termination is defective, non-coding transcripts may be extended and interfere with the expression of nearby genes [115, 116]. For example, mutation of a specific phosphosite in *SEN1* results in readthrough transcription of a long non-coding RNA that interferes with transcription of *ZAP1*, a gene that encodes a master regulator of zinc homeostasis [115]. Extension of a non-coding RNA through the promoter region of *ZAP1* decreased the amount of RNA polymerase II at the promoter, as well as the amount of *ZAP1* RNA. We propose a similar phenomenon is happening at the *IME1* promoter in the *sen1-md* diploid. Failure to terminate some of the *IRT2* transcripts would result in transcription proceeding through the *IRT1* part of the promoter, thereby establishing repressive chromatin that would decrease the amount of *IME1* transcription (Fig 8). Thus, it may take longer to achieve the amount of Ime1 needed to initiate expression of the early genes.

### *SEN1* functions prior to prophase I to prevent *SPO11*-independent DSBs

During S phase in vegetatively growing cells, R-loops accumulate at places where replication collides head-on with transcription (TRCs) [6, 9, 117–121]. When TRCs occur, positive supercoils accumulate between the replication forks and transcription machinery, stalling both processes. The transcribed RNAs may then base pair with the template strand of the DNA to make R-loops [120]. Sen1 associates with the replisome and helps prevent R-loops from forming at TRCs both by displacing RNA Pol II as well as coordinating the activities of topoisomerases I and II to relieve topological stress [117, 118, 120, 121].

R-loops also accumulate at TRCs during premeiotic S phase [31]. When the precise genomic locations of R-loops were mapped in an *rnh1Δ rnh201Δ* SK1 diploid after transfer to Spo medium, an increase in R-loops at TRCs was observed over WT at the 2 hour timepoint when premeiotic S was occurring [31, 122]. No increase in R-loop signal was detected when the orientation of a specific TRC was changed such that replication and transcription were co-directional. The R-loops at premeiotic TRCs are distinct from the DNA:RNA hybrids formed at Spo11-generated DSBs in that they do not co-localize with Spo11 hotspots and their levels are reduced at later timepoints when chromosomes have fully synapsed [31]. Interestingly deletion of *SPO11* increased the frequency of R-loops at the premeiotic TRCs and these R-loops exhibited a similar distribution as the R-loops observed in the *rnh1Δ rnh201Δ* diploid [31].

Several observations support the idea that *SEN1* also plays a role in preventing/removing R-loops at TRCs during premeiotic S phase for the following reasons. (1) *sen1-md* diploids exhibited a severe sporulation defect, in part because some cells arrested in prophase I due to unrepaired DSBs triggering the MRC. If the *sen1-md* sporulation defect was due solely to unrepaired Spo11 DSBs, then *spo11Δ* should have rescued sporulation in the *sen1-md* diploid, but this was not the case. (2) The MRC provides time for DSBs to get repaired, as eliminating the MRC in the *sen1- md* background using either *NDT80-mid* or *mek1Δ* increased the number of fragmented nuclei without improving sporulation. In both cases the sporulation defects were rescued by *PREC8-SEN1*. In the *sen1-md spo11Δ* strain, the MRC was abrogated as expected, with an increase in MI cells compared to *sen1-md*. However, many cells that completed MI did not proceed through MII, perhaps because DSBs triggered the DNA damage checkpoint [76]. Most of the cells that progressed through MII exhibited fragmented nuclei, demonstrating the presence of *SPO11*-independent DSBs. (3) The fact that *PREC8-SEN1* only weakly rescued the sporulation defect of *sen1-md spo11Δ*, while *PSEN1-SEN1* rescued completely, argues that the timing of *PREC8-SEN1* expression was too late in the *sen1-md spo11Δ* diploid to effectively complement *sen1-md*. In contrast, *PREC8-SEN1* expression was not too late to complement the failure to repair Spo11- generated DSBs in *NDT80-mid sen1-md* or *mek1Δ sen1-md* strains. These results suggest the *SPO11*-independent DSBs arise at an earlier time than *SPO11*-dependent prophase I DSBs. (4) DNA:RNA hybrid foci were increased after 2 hours in Spo medium in *sen1-md* compared to both WT at 2 hours and *sen1-md* at 0 hours. This phenotype was exacerbated in the absence of *SPO11*. The latter finding is informative because *spo11Δ* diploids do not make the prophase I DSBs that are used for recombination and therefore the S9.6 foci in the *sen1-md spo11Δ* cannot be attributed to DNA:RNA hybrids at resected DSB ends.

We propose that the *SPO11*-independent DSBs in the *sen1-md spo11Δ* diploid arise from R-loops that form during premeiotic S phase and that *SEN1* functions to prevent/eliminate R loops at premeiotic TRCs. Sen1 could help prevent R-loop formation by removing RNA pol II from TRCs or coordinating topoisomerase I and II activity as it does during vegetative S phase. However, *SEN1* is not sufficient to completely prevent R-loop formation at TRCs, otherwise the R-loop frequency at TRCs would not increase in *rnh1Δ rnh201Δ* diploids where *SEN1* is functional [31]. Therefore, it is also possible that Sen1 may disassemble R-loops at premeiotic TRCs using its helicase activity. The multi-functional roles that *SEN1* may play at premeiotic TRCs could explain why *sen1-md* is more defective in sporulation and spore viability than the *rnh1Δ rnh201Δ* mutant, since RNase H1/2 can only remove R-loops but not prevent their formation [17, 31].

An interesting question is what role *SPO11* is playing at TRCs in premeiotic S phase. During WT meiosis, Spo11 is recruited to chromosome axes where it generates prophase I DSBs after DNA replication [123–125]. During premeiotic S phase, Spo11 may generate DSBs to relieve the topological constraints of the positive supercoils on unreplicated DNA, which would be a novel function for this highly conserved endonuclease.

The fact that progression through MI in *NDT80-mid sen1-md*, *mek1Δ sen1-md* and *sen1-md spo11Δ* diploids was similar indicates that *SPO11*-independent DSBs do not trigger the MRC. Spo11-generated DSBs occur on the chromosome axes, after which the Mec1/Tel1 checkpoint kinases phosphorylate Hop1 [48]. Mek1 binds to phosphorylated Hop1 via its FHA domain and then activates itself by phosphorylation of the Mek1 activation loop in *trans* [50, 51, 126]. We propose that the *SPO11*-independent DSBs arise from processing of persistent R-loops formed at TRCs during premeiotic S phase and thus are likely not forming on chromosome axes, in which case Mek1 (and therefore the MRC) would not be activated [50, 51, 126]. A similar phenomenon was observed in *C. elegans* mutants lacking RNase H activity where R-loops accumulated in the germ line [26]. An increased frequency of DSBs was observed in *rnh-1.0 rnh-2; spo11* worms compared to *spo11* alone and these DSBs failed to trigger the DNA damage checkpoint [26]. Therefore it is possible that a non-canonical function for Spo11 in regulating R-loops during meiosis may be conserved.

### *SEN1* promotes the repair of *SPO11* DSBs during prophase I

Several studies have shown that DNA:RNA hybrids can form on the resected ends of DSBs in a variety of species both in vegetative and meiotic prophase I cells [13, 14, 16, 17, 26, 31, 33, 34]. In budding yeast meiosis, DNA:RNA hybrids that accumulated at the resected ends of DSBs in an *rnh1Δ rnh201Δ hpr1Δ* diploid impaired recombinase binding and interfered with DNA repair [17].

Our work indicates that *SEN1* also functions to promote repair of *SPO11* DSBs.

Meiotic progression in *sen1-md* was delayed due to the MRC that is triggered by Spo11- generated DSBs [50–53]. Furthermore, when *sen1-md* diploids were allowed to progress through meiosis without the MRC-mediated delay, nuclei were fragmented, indicating the presence of broken chromosomes. DNA:RNA hybrid foci were elevated in *sen1-md* strains throughout prophase I. Interestingly, COs were reduced only by 25% in the *sen1-md* diploid and 40% in *rnh1Δ rnh201Δ hpr1Δ* diploid (Fig 4) [17], indicating that only a subset of DSBs was not repaired. This may be due to redundancy between RNase H1/2 and Sen1 in removing the DNA:RNA hybrids at the ends of DSBs, although the *rnh1Δ rnh201Δ sen1-md* diploid was too sick to directly test this idea. Another possibility is that Sen1 and RNase H act on qualitatively different DSBs. Both COs and NCOs were reduced in the *rnh1Δ rnh201Δ hpr1Δ* diploid, while only COs were decreased by *sen1-md* (Fig 4) [17]. Interestingly, chromosome synapsis is defective in the *sen1-md* mutant, despite a relatively small reduction in crossovers at the *HIS4LEU2* hotspot. The ZMM crossover pathway is required for chromosome synapsis [64, 84, 88]. An intriguing idea is that Sen1 specifically removes DNA:RNA hybrids from DSB ends that are destined to be repaired by the ZMM pathway. In this case, most of the COs observed in the *sen1-md* mutant would be generated by the action of structure-selective nucleases such as Mus81-Mms4 that do not contribute to synapsis. RNase H may instead remove DNA:RNA hybrids from DSBs that are processed by other pathways to form either COs or NCOs. Further work is necessary to test this idea.

## ACKNOWLEDGEMENTS

We are grateful to Aaron Neiman for comments on the manuscript. Many thanks to Scott Keeney, Neil Hunter, Raphäelle Laureau, Nicola Silva and members of the Hollingsworth lab for helpful discussions. Craig Chen provided technical help. Neil Hunter, Aaron Neiman, and Liangran Zhang provided strains and/or plasmids.

## METHODS

### Strains and media

All the strains used in this study were derived from the SK1 background. Genotypes of each strain are listed in S1 Table. Growth media are described in [127]. Sporulation (Spo) medium consisted of 2% potassium acetate.

Gene deletions were made using polymerase chain reaction (PCR) generated fragments to replace open reading frames (ORFs) with either *kanMX6* (confers G418 resistance), *natMX4* [confers nourseothricin (NAT) resistance], or *hphMX4* (confers Hygromycin B resistance) using the plasmids pFA6a-kanMX6, p4339, or pAG32, respectively. Deletions were verified by PCR using a forward primer upstream of the deleted ORF and a reverse primer within the drug marker. In addition, the absence of the WT gene was confirmed by failure to detect a PCR product using the same forward primer and a reverse primer in the ORF. For the *ndt80Δ::hphMX4* haploid parents of AM6697, flanking primers generated different sized PCR products for the *NDT80* and *ndt80Δ::hphMX4* alleles.

To make *sen1-md* strains, a 2.5 kb fragment was amplified from pMJ787 containing *kanMX6* 133 bp upstream of *PCLB2-3xHA* . This fragment was designed with homologies such that it replaces the 50 bp immediately upstream of the *SEN1* ATG with the *CLB2* promoter. G418^R^ haploid transformants of both mating types were screened by colony PCR to confirm the insertion of the fragment directly upstream of *SEN1* using a forward primer located 814 bp upstream of *SEN1* and a reverse primer 740 bp downstream of the *SEN1* start codon.

The *PREC8-SEN1* and *PREC8-sen1-ΔN* plasmids, pNH410 and pBG27, respectively, were digested with PshAI to target integration to codon 1811 of *SEN1* and transformed into the *sen1-md* diploid, NH2667. Integration of the plasmids was confirmed by complementation of the sen1-md sporulation defect and/or immunoblots probed with α-Sen1 antibodies.

The *sen1-md NDT80-mid* diploid, NH2667::pNH317^2^ was created by transforming the haploid parents of NH2667 with the *URA3 NDT80-mid* integrating plasmid, pNH317, digested with SnaBI to target integration upstream of the *NDT80* ORF (note that *NDT80-mid* is dominant to *NDT80*)[52]. Proper plasmid integration was by PCR with a forward M13 primer present in the vector sequence with a reverse primer 1.9 kb upstream of the *NDT80-mid* ORF.

### Plasmids

The genotypes of plasmids used are listed in S2 Table. Any sequences amplified by PCR were sequenced in the final plasmids by the Stony Brook University DNA Sequencing Facility or by Plasmidsaurus (https://www.plasmidsaurus.com/).

The *SEN1* gene was fused to the *REC8* promoter in pNH257 to make pNH410 using the NEBuilder HiFi DNA Assembly Master Mix (hereafter referred to as Gibson assembly or GA) (New England Biolabs, Cat.# E2621). A 2μ *SEN1* plasmid containing a genomic fragment from chromosome XII with the coordinates 989631-1000863 was used as the template for PCR [128]. The 6696 bp *SEN1* ORF was amplified as three separate fragments of approximately 2 kb. The SEN1a fragment contains 25 bp of homology to the BamHI cut site of pNH257 (immediately downstream of *PREC8*) fused to *SEN1* codons 1-790. SEN1b contains codons 783-1555 and SEN1c contains codons 1547-2231 along with 210 bp of 3’ untranslated sequence and 25 bp of homology to the EcoRI cut site of pNH257. The three PCR fragments were incubated with BamHI/EcoRI- digested pNH257 in a GA reaction following the manufacturer’s instructions. The *PREC8- sen1-ΔN* allele in pBG27 lacks the first 1003 codons of *SEN1*. pNH410 was used as the template to generate a 1691 bp PCR fragment containing homology to the EcoRI site on pNH257 along with an ATG and *SEN1* codons 1004-1555. This fragment was then joined to the SEN1c fragment and ligated into pNH257 using GA as described for pNH410. The *PSEN1-SEN1* plasmid, pBG28, was constructed using a SEN1a fragment that had 25 bp of homology to the NotI site in pRS306 fused to a 2.8 kb sequence containing 0.8 kb located immediately upstream of the *SEN1* ORF, as well as the first 791 codons of the gene. The SEN1b fragment was the same as for pNH410, while the SEN1c fragment contained the same fragment of *SEN1* with 192 bp 3’ untranslated sequence and 3’ homology to the SalI site in pRS306.

### Timecourses

Liquid sporulation was performed as described in [127] with the following changes. Diploids were streaked out from the -80°C freezer onto YPD medium supplemented with complete powder [127] and incubated at 30°C for 2-3 days. Single colonies were inoculated into 8 mL YPD and incubated at 30°C for 24 h. 1.0 mL and 1.7 mL of inoculum were then diluted into 200 mL YPA in two 2 L flasks and incubated at 240 RPM for 18 hours at 30°C. The optical density at 660nm (OD660) of a 2-fold dilution in YPA of the culture was read using a SPECTRONIC 200 spectrophotometer. Diluted cultures with OD660 readings between 1.6 and 2.3 were pelleted, washed with water and resuspended in the volume of Spo medium required for a cell density of 3 x 10^7^ cells/mL in 2 L flasks. The cells were incubated in a 30°C shaker rotating at 250 RPM. At each timepoint, cells were fixed with a 1/10 volume of 37% formaldehyde for 4’,5-diamidino-2- phenylindole (DAPI) staining. Formaldehyde-fixed cells were processed for whole cell immunofluorescence as described in (Ziesel *et al.* 2022). For protein and RNA sequencing samples, 5 ml sporulating culture were pelleted in a 15 ml conical tube, the supernatant was discarded, and the cell pellets stored at -80°C. For DNA physical analysis, 10 mL culture was added to 1 mL 0.5M EDTA and 10 mL 100% ethanol and stored at -20°C. For nuclear spreads, 6.7 mL of sporulating culture were immediately processed as described in [129].

### Nuclear Spreads

#### For localization of meiotic axis and central region proteins of the synaptonemal complex (**Fig 5**)

Strains homozygous for *ndt80::ΔhphMX4* were cultured overnight in BYTA (1% yeast extract, 2% bactopeptone, 1% potassium acetate, 50 mM potassium phthalate) then sporulated in 1% potassium acetate for 5 hours. Nuclei from these cells were surface-spread on glass slides and imaged as described in [130]. Affinity purified rabbit anti-Zip1 (1:150), mouse anti-Gmc2 (1:800), and rabbit anti-Red1 (1:200) [59].

Secondary antibodies conjugated with Alexa Fluor dyes (Jackson ImmunoResearch) were used at a 1:200 dilution. Microscopy and image processing were performed using a Deltavision RT imaging system (General Electric) adapted to an Olympus (IX71) microscope.

#### For localization of DNA:RNA hybrids (**Fig 7**)

Meiotic cultures were spheroplasted and their nuclei spread and immunostained as described in [129]. Slides treated with 5U of RNase H (New England Bioscience M0297L) were incubated at 37°C for 1h in a moist slide box after blocking and then washed with 1xTBS prior to incubation with antibodies. All immunostains were incubated with antibodies at 4°C in a moist slide box. Slides were first stained for DNA:RNA hybrids with mouse anti-DNA:RNA hybrids (S9.6) (Kerafast ENH001) diluted 1:1000 in 1% BSA and incubated overnight. After washing slides as described, goat α-mouse IgG-Alexa 594 (Fisher A32742) secondary antibody was diluted 1:1000 in 1% BSA was applied to slides and incubated for two hours. Micrographs were taken with a ZEISS Axio Imager.Z2 with a Zeiss Plan-Apochromat 100X objective and analyzed using ZEN 3.5 software.

#### Physical Analysis of DSBs and Repair Products

Genomic DNA was isolated from ethanol-fixed sporulation samples using the MasterPure Yeast DNA Purifiation Kit (Biosearch Technologies MPY80200) and quantified with the Qubit 1x dsDNA HS Assay Kit (Invitrogen Q33231). Digestion of samples, gel electrophoresis, Southern blotting, and hybridization with radiolabeled probe for analysis of the *HIS4-LEU2* hotspot were performed as described in (Owens et al. 2018) with the following changes: the blotted membrane was incubated in prehybridization solution (0.5% SDS, 0.5 mg/mL sheared salmon sperm DNA, 6xSSC (29.22 mg/mL sodium chloride, 14.7mg/mL sodium citrate, HCl to pH 7.0)) at 68°C for 2 hours. ^32^P-labeled probe with at least 1x10^6^ counts per minute was added to hybridization solution (0.5% SDS, 0.5 mg/mL sheared salmon sperm DNA, 6xSSC, 0.1 mg/mL dextran sulfate) and incubated with the membrane for 16h at 68°C. The membrane was washed at 68°C three times for 30 minutes with ∼40mL of Wash Buffer 1 (2x SSC, 0.1% SDS) and then twice with Wash Buffer 2 (0.2x SSC, 0.1% SDS). It was then washed at room temperature for 5 minutes each with Wash Buffer 2 and Wash Buffer 3 (0.1x SSC). The blots were imaged using a FujiFilm FLA-7000 phosphoimager and Multi-Gage software.

#### Antibodies

The sources and conditions used for various antibodies are listed in S3 Table. The Sen1 antibody was generated by Covance Research Products (now Labcorp Drug Development) in a guinea pig using the peptide Ac-CFSDDVSFIPRNDEPEIK-amide (amino acids 2002-2019).

#### RNA sequencing analysis

After growth in YPA, the *SEN1* and *sen1-md* diploids were transferred to Spo medium at a density of 3 X 10^7^ cells/ml and 5 ml aliquots were taken at various timepoints, pelleted and frozen at -80°C. The cell pellets were sent to GENEWIZ (now Azenta) for RNA extraction and polyA RNA enrichment. The kit used to prepare the libraries was the NEBNext Ultra RNA Library Prep, PolyA (New England Biolabs). The adapter sequences used to generate the RNA sequencing data were: Read1: AGATCGGAAGAGCACACGTCTGAACTCCAGTCAC Read2: AGATCGGAAGAGCGTCGTGTAGGGAAAGAGTGTA The RNA sequencing data were mapped to the yeast genome using Bowtie 1.1.2 and custom Python scripts with Yeast PacBio 2016 as the reference genome (https://yjx1217.github.io/Yeast_PacBio_2016/data/)[131] (Langmead B et al. doi: 10.1186/gb-2009-10-3-r25). Fragments/kb transcript/million (fpkm) values were determined using TopHat2 (https://doi.org/10.1186/gb-2013-14-4-r36), which were then used to calculate the standard scores (Z-scores) for each gene using the formula (fpkmn - fpkmave)/SD, where n = timepoint, fpkmave = average of the fpkm values for each timepoint and SD is the standard deviation. Genes were clustered based on their pattern of Z values using Clustal 3.0 and then visualized as a heat map plotted by JavaTree. The *sen1-md* strain was not weighted and these genes were “passengers” whose location was set by the WT cluster.

## Supporting Information

**S1_Data.** Contains the data and calculations used for all of the numerical data presented in Figures. (XLXS)

**S1 Fig.**
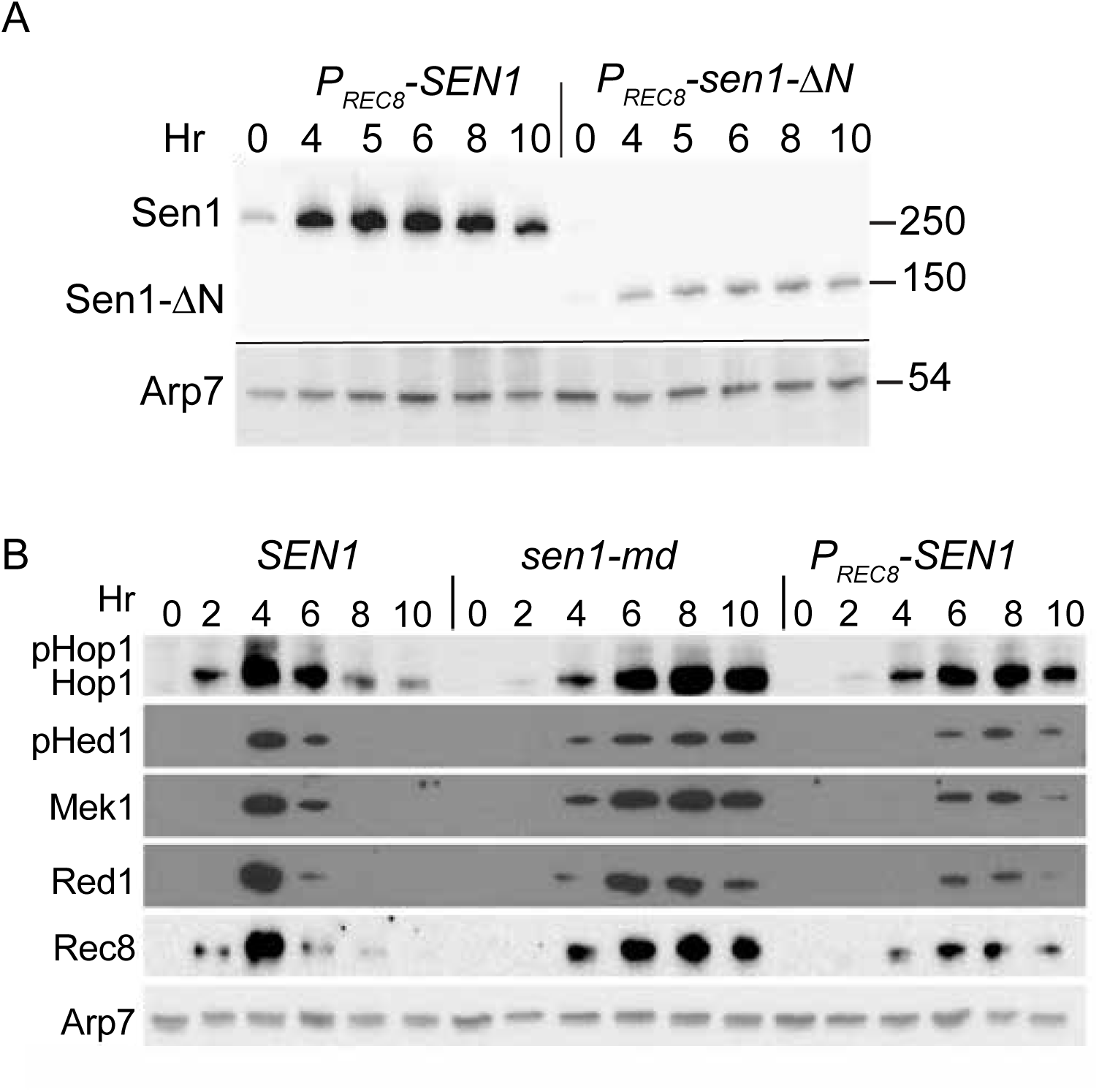
Immunoblots using a newly developed α-Sen1 antibody. The *sen1-md* diploid, NH2667, was transformed with an integrating plasmid containing either *PREC8- SEN1* (pNH410) or *PREC8-sen1-ΔN* (pBG27) and induced to undergo meiosis. Protein samples from the indicated timepoints were probed with α-Sen1 antibodies. α-Arp7 antibodies were used to detect Arp7 as a loading control. Numbers on the right indicate the positions of molecular weight markers in kiloDaltons. The black line indicates that the same samples were run on two different gels and probed with different antibodies. “Hr” refers to hours in Spo medium. (B) Timing of expression of various meiosis-specific proteins. Protein extracts from meiotic timecourses using *SEN1*, *sen1-md* and *PREC8- SEN1* strains were probed with antibodies against the indicated proteins. “pHop” and “pHed1” indicate phosphorylated Hop1 and Hed1, respectively.

**S2 Fig.**
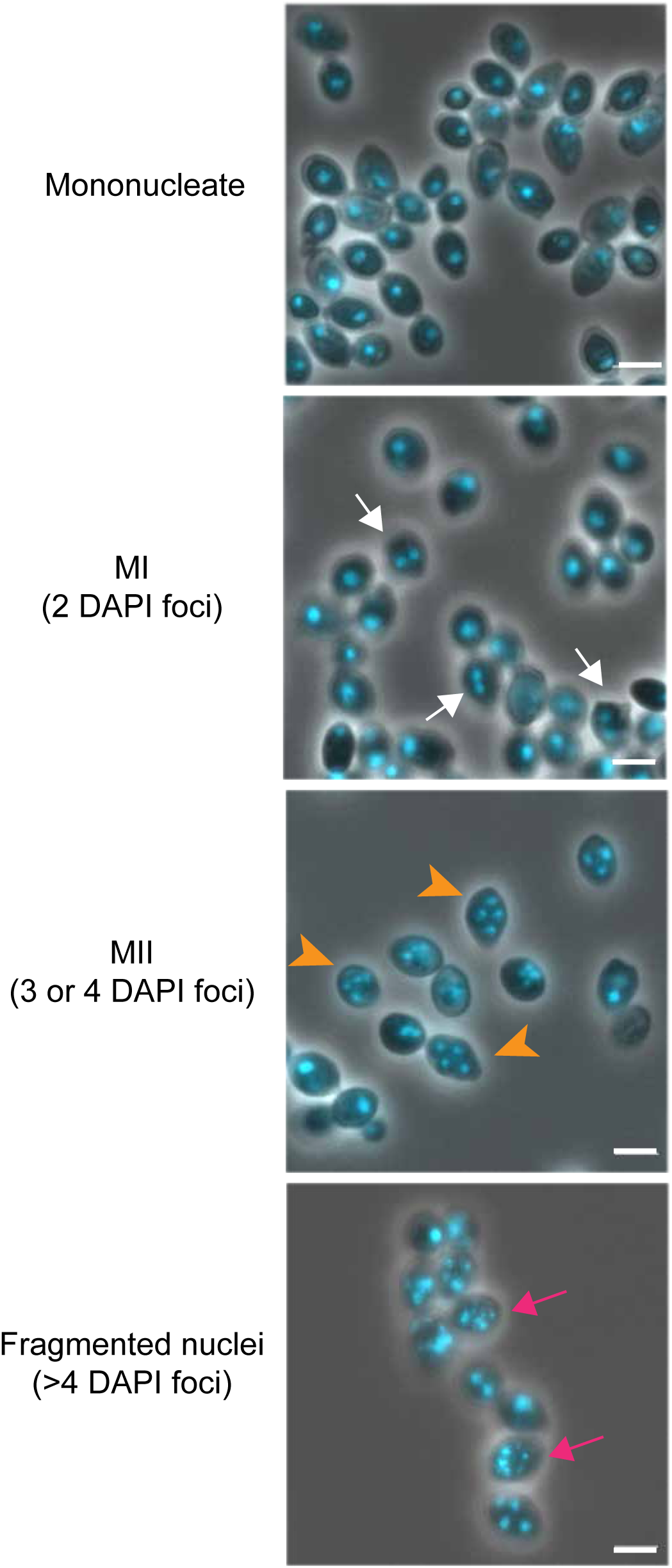
Monitoring meiotic progression and nuclear fragmentation by fluorescent staining of the DNA. Cells were fixed with 3.7% formaldehyde, stained with DAPI and examined by fluorescence microscopy. Mononucleate cells are either vegetative cells or meiotic cells prior to anaphase I. White arrows indicate binucleate MI cells. Orange arrowheads indicate tetranucleate MII cells and magenta arrows indicate cells with fragmented nuclei that contain >4 DAPI foci. The first three images are WT cells (NH716) while the last image is from a *mek1Δ sen1-md* diploid (NH2669)

**S1 Table.**
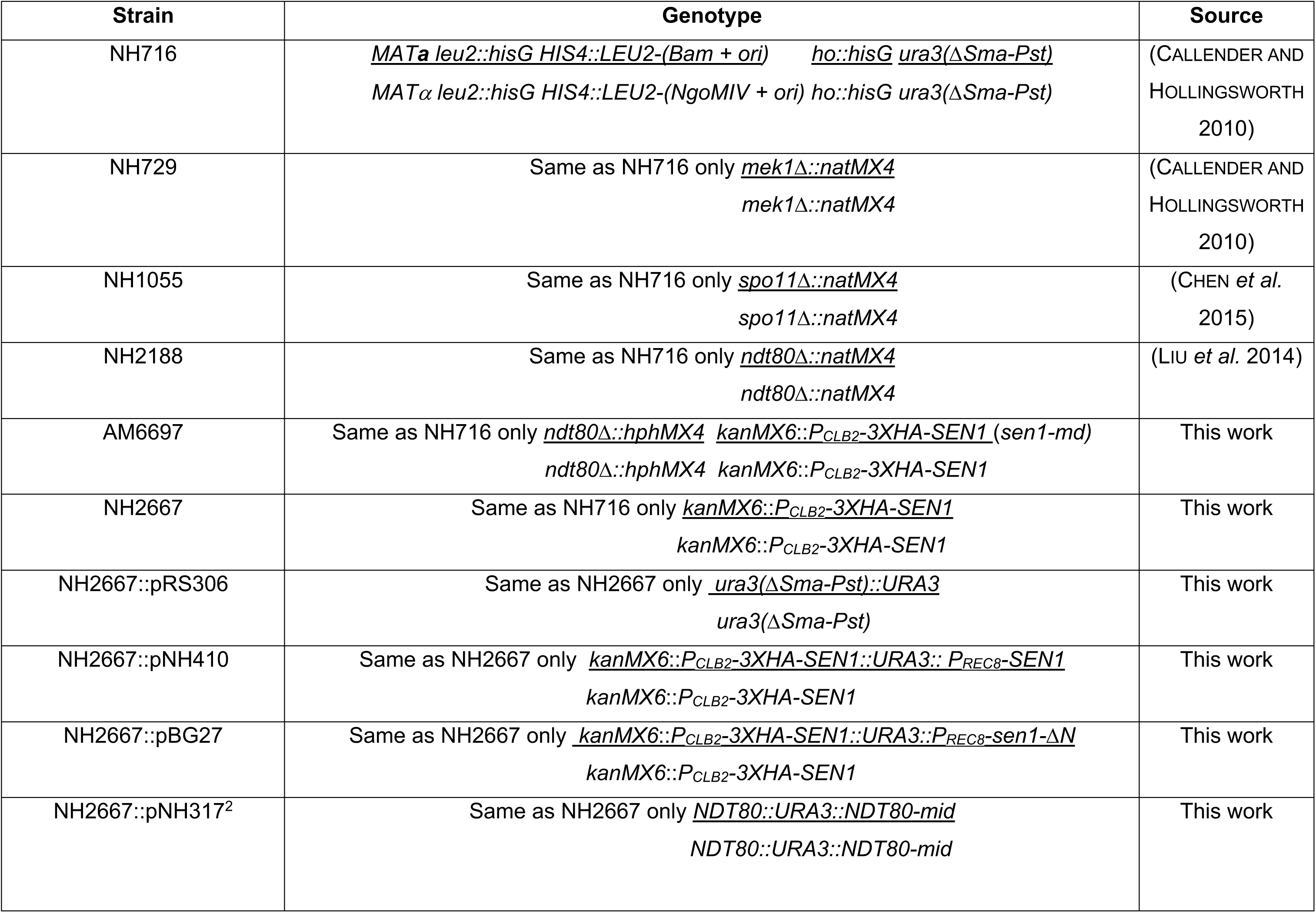
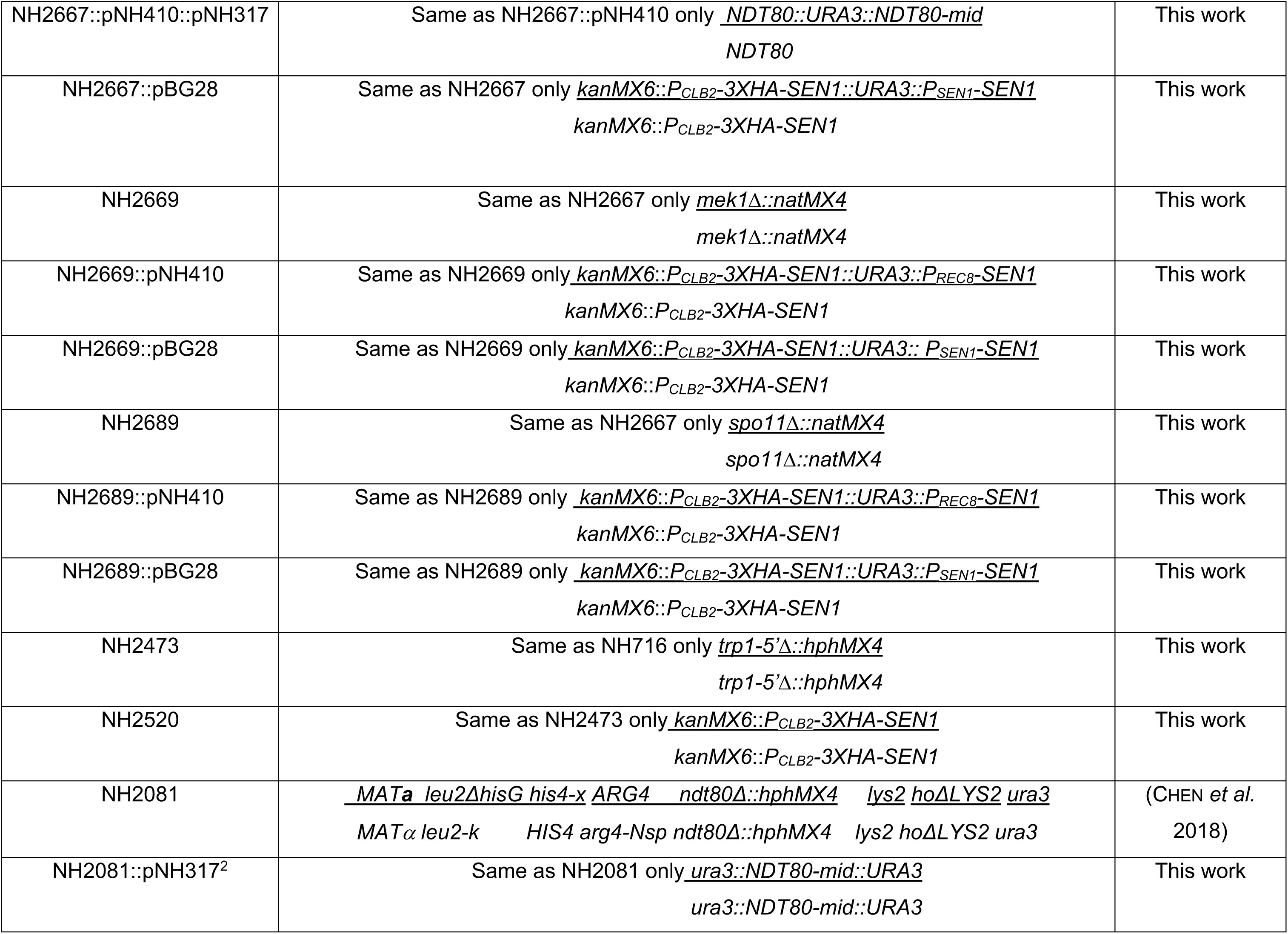
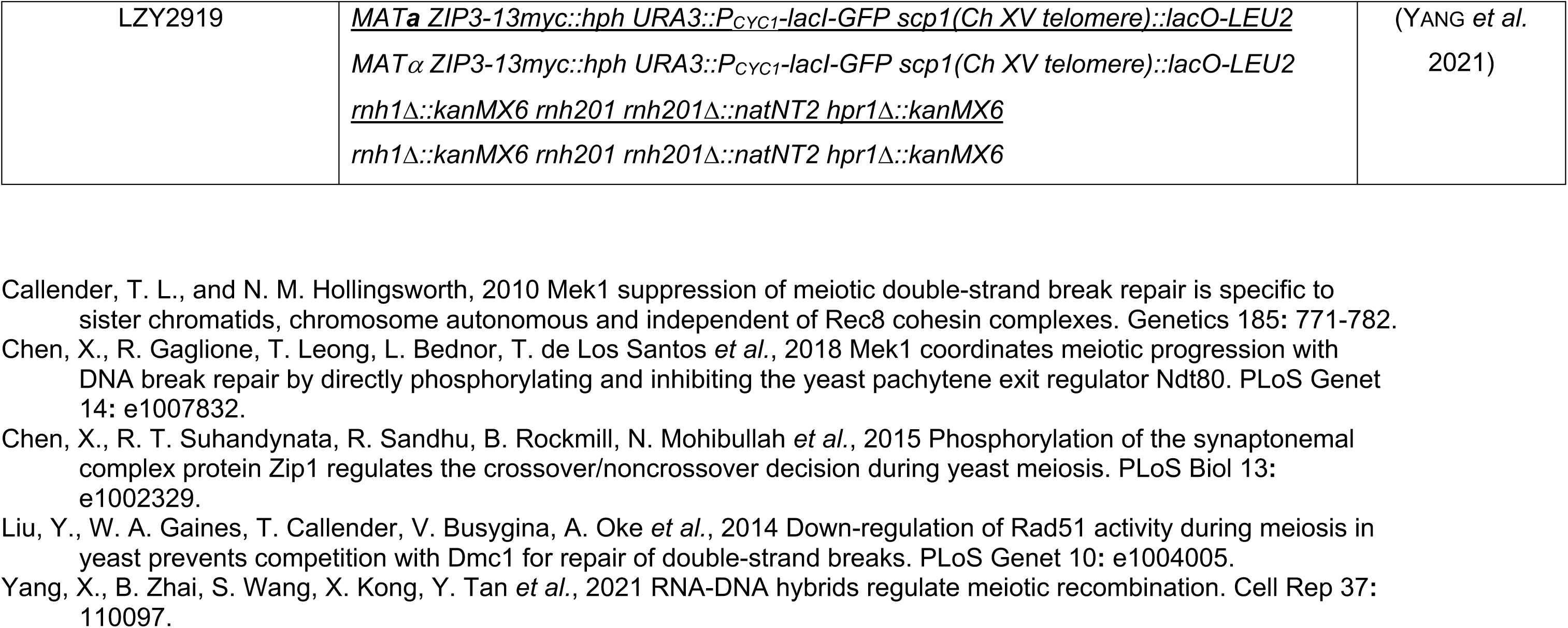
Saccharomyces cerevisiae strains.

**S2 Table.** Plasmids.

| **Plasmid name** | **Relevant yeast genotype** | **Source** |
| --- | --- | --- |
| pRS306 | *URA3* | (PARENT *et al.* 1985) |
| p4339 | *natMX4* | (GOLDSTEIN AND McCUSKER 1999) |
| pAG32 | *hphMX4* | (GOLDSTEIN AND McCUSKER 1999) |
| pFA6a-kanMX6 | *kanMX6* | (LONGTINE *et al.* 1998) |
| pMJ787 | *kanMX6::P<sub>CLB2</sub>-3xHA* | (LEE AND AMON 2003) |
| 2μ *SEN1* | 2μ *LEU2 SEN1* | (JONES *et al.* 2008) |
| pNH257 | *P<sub>REC8</sub> URA3* | (ZIESEL *et al.* 2022) |
| pNH410 | *P<sub>REC8</sub>-SEN1 URA3* | This work |
| pBG27 | *P<sub>REC8</sub>-sen1<sup>1004-2231</sup> (ΔN) URA3* | This work |
| pNH317 | *NDT80-mid URA3* | (CHEN *et al.* 2018) |
| pBG28 | *P<sub>SEN1</sub>-SEN1 URA3* | This work |

**S3 Table.** Antibodies.

| Antigen | Primary | Purpose | Dilution | Source | Secondary | Dilution | Source |
| --- | --- | --- | --- | --- | --- | --- | --- |
| Arp7 | $\alpha$ -Arp7 | Immunoblot | 1:50,000 | Santa Cruz sc-8961 | α-Goat | 1:15,000 | Santa Cruz sc-2354 |
| pHed1 | $\alpha$ -pT40 Hed1 | Immunoblot | 1:20,000 | N. M Hollingsworth | $\alpha$ -Rabbit | 1:10,000 | Invitrogen 31460 |
| Hop1 | $\alpha$ -Hop1 | Immunoblot | 1:10,000 | N. M Hollingsworth | $\alpha$ -Rabbit | 1:10,000 | Invitrogen 31460 |
| Mek1 | $\alpha$ -Mek1 | Immunoblot | 1:10,000 | N. M. Hollingsworth | $\alpha$ -Guinea Pig | 1:10,000 | Santa Cruz sc-2903 |
| Rec8 | $\alpha$ -Rec8 | Immunoblot | 1:50,000 | N. M. Hollingsworth | $\alpha$ -Guinea Pig | 1:10,000 | Santa Cruz sc-2903 |
| Red1 | $\alpha$ -GST-Red1 | Immunoblot | 1:10,000 | N. M Hollingsworth | $\alpha$ -Rabbit | 1:10,000 | Invitrogen 31460 |
| Red1 | $\alpha$ -GST-Red1 | Immunostaining – Figure 1 | 1:200 | N. M Hollingsworth | $\alpha$ -Rabbit – Alexa 488 | 1:1,000 | Invitrogen A11008 |
| Red1 | $\alpha$ -Red1 | Immunostaining – Figure 5 | 1:200 | G. S. Roeder | $\alpha$ -Rabbit – Alexa 488 | 1:200 | Jackson – 711-545-152 |
| Gmc2 | $\alpha$ -Gmc2 | Immunostaining – Figure 5 | 1:800 | A. J. MacQueen | $\alpha$ -mouse – Alexa 594 or<br>$\alpha$ -mouse – Alexa 488 | 1:200 | Jackson-715-585-150 or<br>715-545-150 |
| Zip1 | $\alpha$ -Zip1 | Immunostaining | 1:150 | A. J. MacQueen | $\alpha$ -Rabbit – Alexa 594 | 1:200 | Jackson – 711-585-152 |
| S9.6 | $\alpha$ -RNA – DNA hybrid | Immunostaining | 1:1000 | Kerafast - ENH001 | $\alpha$ -mouse – Alexa 594 | 1:10,000 | Invitrogen A32742 |
| Sen1 | $\alpha$ -Sen1 | Immunoblot | 1:5,000 | N. M Hollingsworth | $\alpha$ -Guinea Pig | 1:10,000 | Santa Cruz sc-2903 |

## Notes

### Competing Interest Statement

The authors have declared no competing interest.

